# Laminin Receptor Characterization in Acute Myeloid Leukemia: Integrin α7β1 Defines non-Leukemic Stem Cells with Migratory Potential

**DOI:** 10.1101/2024.03.29.587290

**Authors:** Elsa Görsch, Marlon Arnone, Maksim Klimiankou, Jan Weller, Saskia Rudat, Gerd Klein, Claudia Lengerke

**Affiliations:** University Hospital Tübingen, Department for Internal Medicine II, University of Tübingen, Tübingen, Germany; German Cancer Consortium (DKTK), partner site Tübingen, a partnership between DKFZ and University Hospital Tübingen, Germany

## Abstract

Interactions with the bone marrow (BM) niche are crucial for promoting self-renewal and survival of acute myeloid leukemia (AML) cells. Consequently, AML cells express a variety of surface receptors to engage with BM niche cells and extracellular matrix proteins, including laminins. Despite the association of laminin receptor expression with stemness in healthy hematopoiesis, the role of laminin receptors in AML remains poorly understood. In this study, we present a comprehensive examination of the laminin receptors integrin α3β1, α6β1, α7β1 and basal cell adhesion molecule (BCAM) in AML. We demonstrate that high mRNA expression of all four laminin receptors correlates with poor overall survival. Notably, integrin α6 and α7 display the highest cell surface presentation among the examined laminin receptors and are higher expressed on AML cells compared to healthy controls. Moreover, our results indicate that integrin α7 expression allows to distinguish between leukemic stem cells (LSC) and non-LSC populations. Specifically, integrin α7 appears to mark non-LSC with enhanced migratory potential. Together, our results confirm the association of high laminin receptor expression with poor prognosis and establish integrin α7 as marker of high migratory non-LSC.

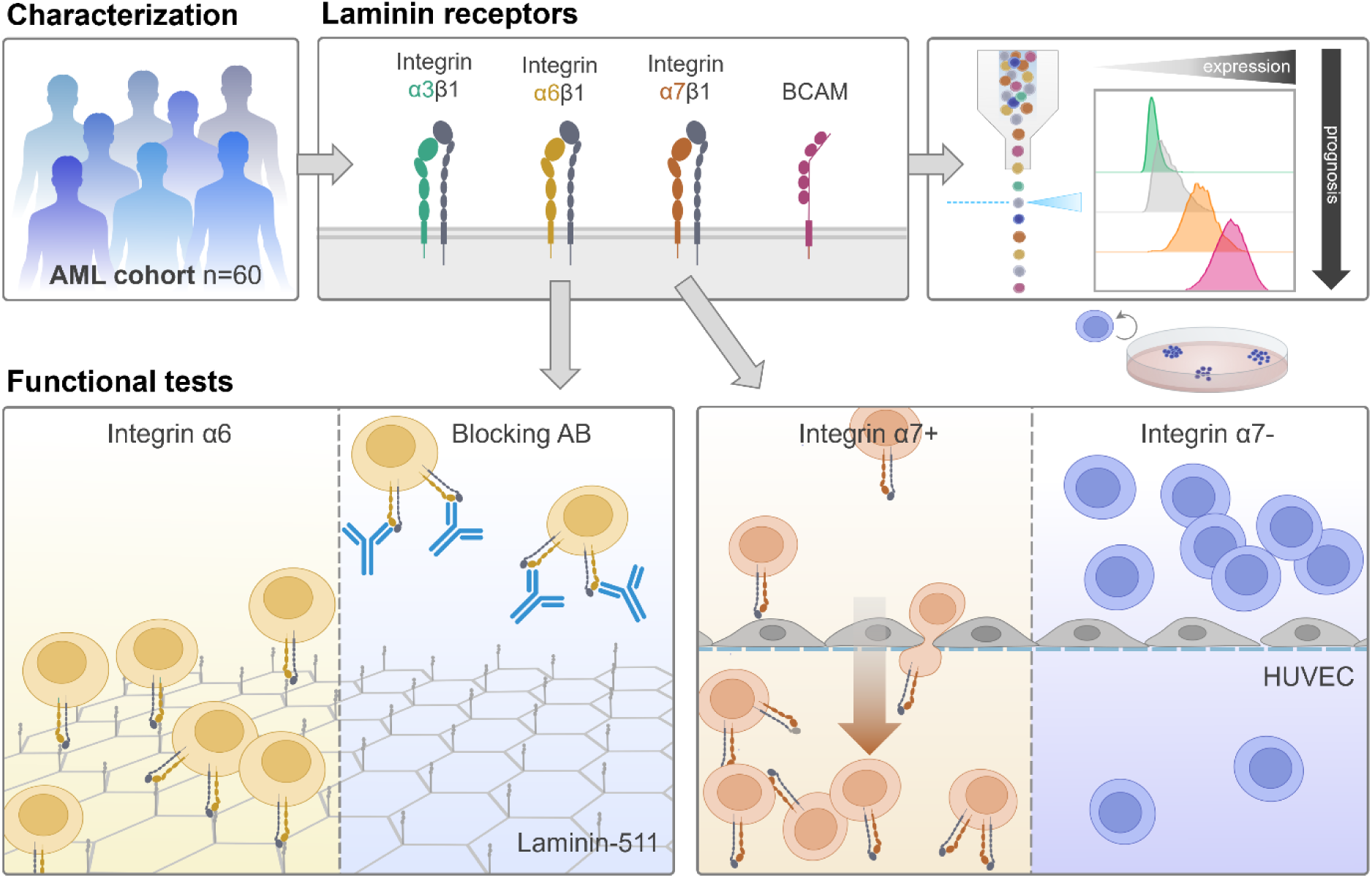

## INTRODUCTION

Acute myeloid leukemia (AML) is driven by leukemic stem cells (LSC) that result from the malignant transformation of healthy hematopoietic stem and progenitor cells (HSPC) upon acquisition of genetic and/or epigenetic lesions. LSC not only initiate leukemia, but also contribute to chemoresistance and relapse (1–3). In contrast, non-LSC are more easily cleared by therapies and are not considered leukemogenic (4).

Like healthy HSPC, LSC depend on interactions with the bone marrow (BM) microenvironment to maintain self-renewal and to promote survival and therapy resistance (5). The concept of the BM niche as a specialized microenvironment enabling hematopoietic stem cell homeostasis was postulated by Raymond Schofield already back in 1978 (6), and numerous follow-up studies on the architecture and composition of the BM niche have detailed its control on hematopoiesis and hematopoietic stem cell fate (7–9). In addition to cellular components, the BM niche further contains a complex extracellular matrix (ECM) (10,11). The ECM provides a structural scaffold and furthermore harbors ECM-bound growth factors influencing cell signaling. Interactions between ECM components and niche cells are mediated by cellular receptors, of which integrins form the largest group. Integrins are heterodimeric transmembrane proteins composed of an α- and β-subunit. 18 different α- and β-subunits are combined to form a total of 24 integrins (12,13). Most integrins contain a β1-subunit, including those with laminin binding specificity, integrin α3β1, α6β1 and α7β1 (14,15).

Integrins bind different components of the ECM, such as collagens, fibronectins or laminins (16,17), and thereby mediate pivotal roles in cell adhesion. In healthy hematopoiesis, laminin receptors are associated with stemness. Integrin α3 (ITGA3, CD49c) has been proposed as a marker for cultured functional long-term HSC (18) and integrin α6 (ITGA6, CD49f) expression may be used in combination with other markers to distinguish long-term multilineage engrafting HSC and hematopoietic progenitors (19). Furthermore, BCAM and integrin α7 have been shown to be expressed by human Lin-CD34+ HSPC (20).

Leukemic stem and progenitor cells are known to reside in and home to BM niches and are considered to outcompete their healthy counterparts (21). However, our understanding of the role of laminin receptors in AML is still rudimental. So far, high expression of integrin α6 has been shown to contribute to drug resistance in EVI1^high^ AML (22) and high integrin α7 expression has been associated with poor prognosis and the occurrence of extramedullary AML (23,24). With this study, we provide a comprehensive characterization of the laminin receptors α3β1, α6β1, α7β1 and BCAM in AML.

## MATERIALS AND METHODS

### Isolation of primary AML, healthy mobilized HSPC donor PBMC and healthy cord blood mononuclear cells

Human cord blood (CB), peripheral blood (PB) and BM samples were obtained from the University Hospitals of Tübingen and Basel with informed consent and approval by the respective Ethics Review Boards (Table 6). Mononuclear cells (MNC) were isolated by density gradient centrifugation. In brief, samples were diluted 1:1 with phosphate-buffered saline (PBS) and MNC were enriched using SepMate-50 tubes (Stem Cell Technologies, 85460) and Pancoll separation solution (PAN Biotech, P04-20150). Red blood cells were lysed using ACK lysis buffer and MNC were viably frozen in RPMI 1640 medium (ThermoFischer, 21875034), 20 % fetal bovine serum (FBS, ThermoFisher, A5256701) and 10 % dimethyl sulfoxide (DMSO) and stored in liquid nitrogen.

### Purification and culture of CD34+ cord blood HSPC

CD34+ HSPC were enriched from cord blood mononuclear cells (MNC) by magnetic selection using anti-CD34 conjugated magnetic beads according to the manufacturer’s instructions (Miltenyi Biotec, 130-100-453). To obtain a sufficient cell number for ImageStream and Western blot analysis, CD34+ HSPC were expanded for 4 days in StemMACS medium (Miltenyi, 130-100-463), 10 % FBS, 50 ng/mL cytokines (FLT3-L, SCF, TPO, G-CSF, IL-3, IL-6), 0.75 µM SR-1 and 0.75 µM UM729.

### Cell culture

AML cell lines (Kasumi-1, MOLM-13, NOMO-1, SKM-1, THP-1) were obtained from DSMZ and human umbilical cord endothelial cells (HUVEC) from ATCC. Cell lines were routinely tested for mycoplasma infection and authenticated using STR assays. AML cell lines were cultured in RPMI 1640 medium (ThermoFischer, 21875034), 10-20 % FBS (ThermoFisher, A5256701), 1% penicillin-streptomycin (ThermoFisher, 15140122) and passaged 1:4 every 3-4 days. HUVEC were seeded at a density of 20 000 cells/cm^2^, cultured in endothelial cell medium (Promocell, C-22010) until they reached 80 % confluence and passaged every 3 days.

### Generation of knockout cell lines

For gene editing, AML cell lines were transfected with ribonucleoprotein (RNP) complexes. To transfect 1x10^6^ cells, 220 pmol sgRNA (IDT) were mixed with 20 pmol SpCas9 nuclease (IDT, 10007807) and incubated for 15 min at RT to allow RNP formation. Subsequently, RNP complexes were mixed with 100 µL nucleofection buffer (Lonza, 197168). Cells were washed 3x with PBS, resuspended in 100 µL nucleofection buffer containing RNP and electroporated according to manufacturer’s instructions using the 4D-Nucleofector (Lonza). Control cells were electroporated with SpCas9 nuclease only.

### Flow cytometry analysis and cell sorting

For phenotyping, 1x10^6^ cells were washed once with FACS buffer (PBS, 5 % FBS, 2 mM EDTA) and subsequent staining steps were performed in FACS buffer using the antibodies depicted in table 1. Cells were acquired with a BD LSR Fortessa flow cytometer (BD Biosciences) and data analysis was performed using FlowJo (version 10.8.0). Dead cells were excluded from the analysis using a viability dye (Thermo Fisher Scientific, L34966) and CD33 was used as a marker for AML blasts. 20x10^6^ cells were stained for cell sorting using CD33 to enrich AML blasts and the marker of interest. Cells were sorted with the sorters BD FACS Aria IIIu (BD Biosciences) and MA900 Multi-Application (Sony) excluding dead cells and CD33-negative cells. For gating strategies see **Supplemental Figure S6**.

**Table 1:**
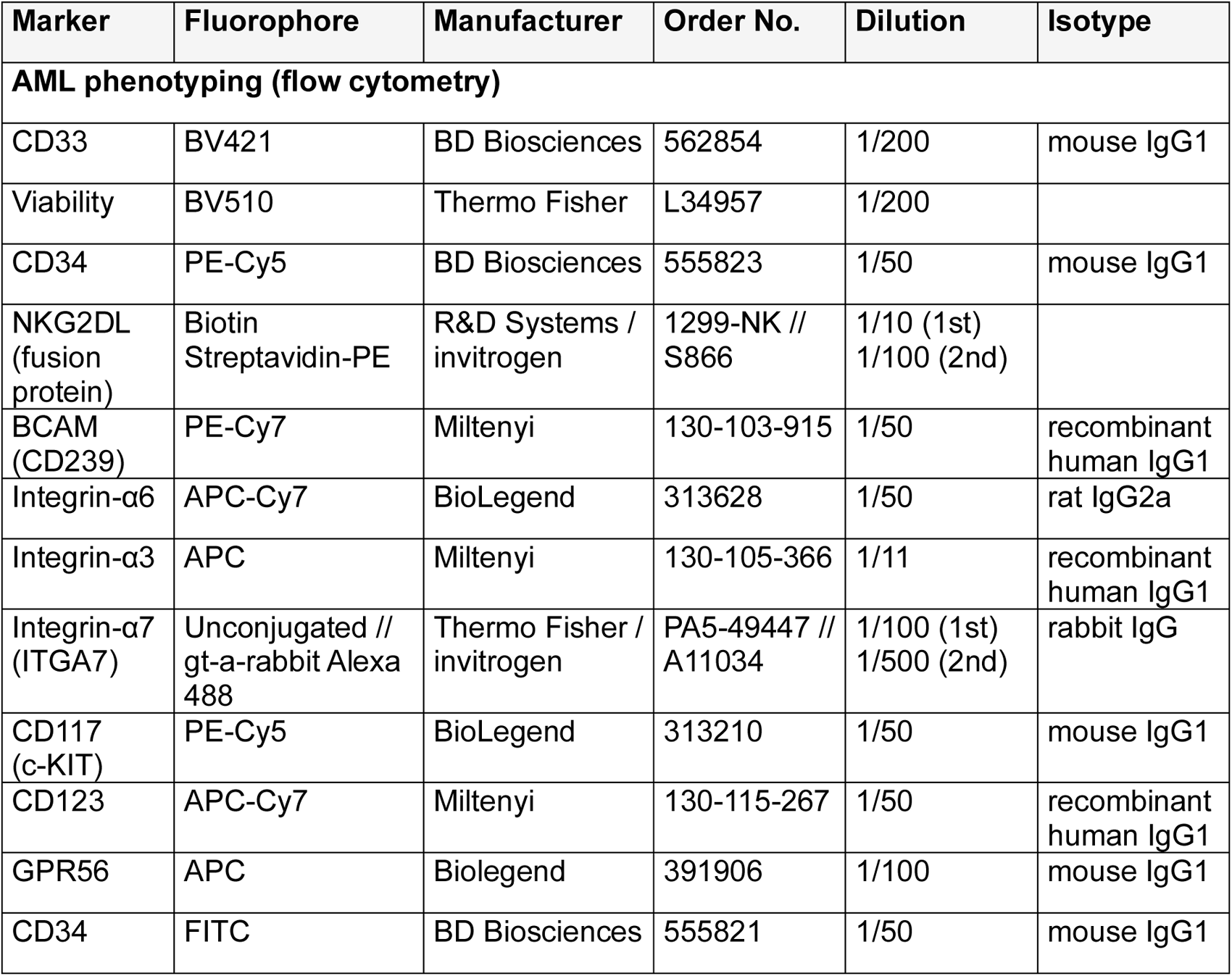
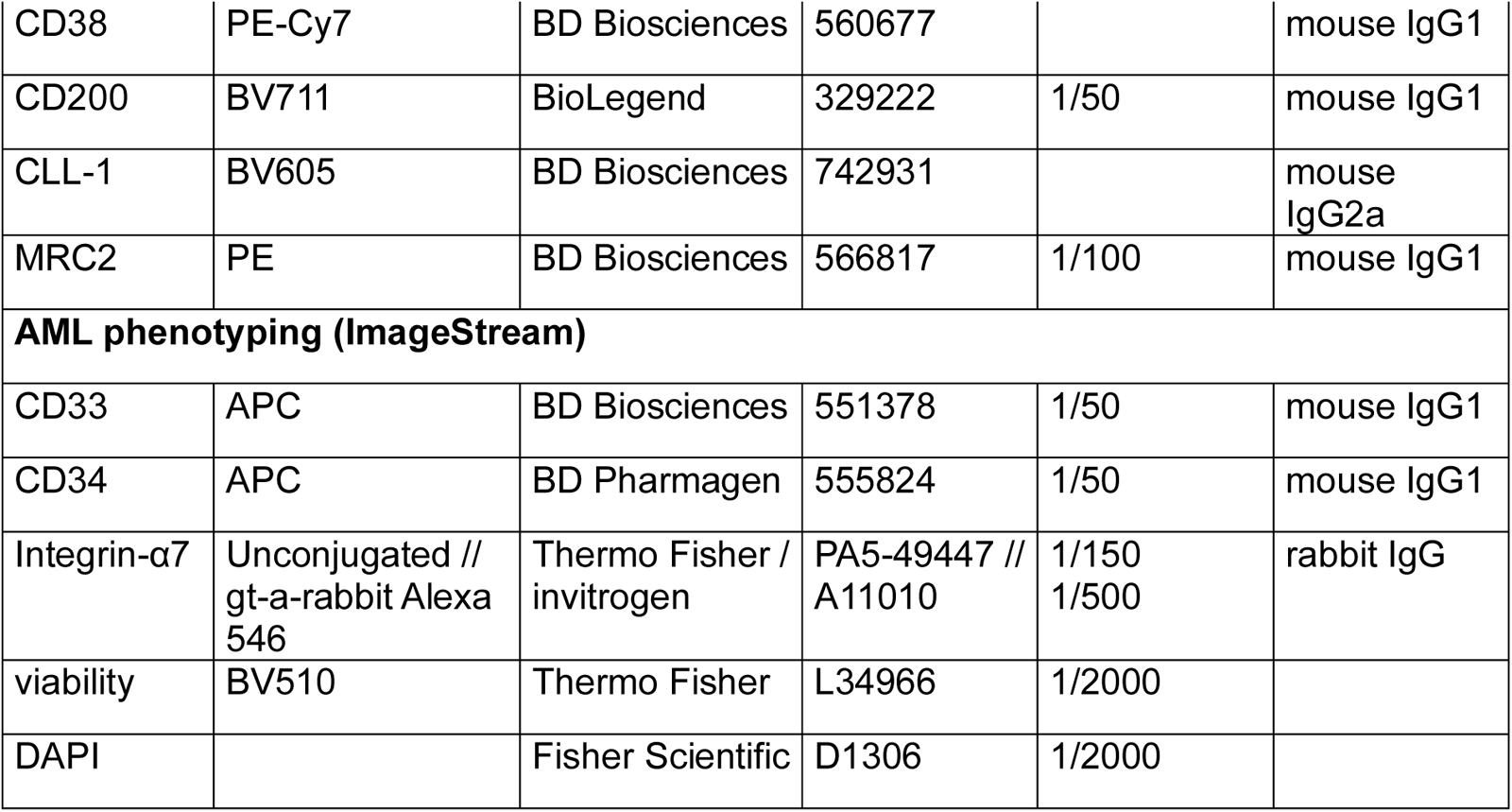
Antibodies for flow cytometry.

| Marker | Fluorophore | Manufacturer | Order No. | Dilution | Isotype |
| --- | --- | --- | --- | --- | --- |
| **AML phenotyping (flow cytometry)** |  |  |  |  |  |
| CD33 | BV421 | BD Biosciences | 562854 | 1/200 | mouse IgG1 |
| Viability | BV510 | Thermo Fisher | L34957 | 1/200 |  |
| CD34 | PE-Cy5 | BD Biosciences | 555823 | 1/50 | mouse IgG1 |
| NKG2DL<br>(fusion protein) | Biotin<br>Streptavidin-PE | R&D Systems /<br>invitrogen | 1299-NK //<br>S866 | 1/10 (1st)<br>1/100 (2nd) |  |
| BCAM<br>(CD239) | PE-Cy7 | Miltenyi | 130-103-915 | 1/50 | recombinant<br>human IgG1 |
| Integrin-α6 | APC-Cy7 | BioLegend | 313628 | 1/50 | rat IgG2a |
| Integrin-α3 | APC | Miltenyi | 130-105-366 | 1/11 | recombinant<br>human IgG1 |
| Integrin-α7<br>(ITGA7) | Unconjugated //<br>gt-a-rabbit Alexa<br>488 | Thermo Fisher /<br>invitrogen | PA5-49447 //<br>A11034 | 1/100 (1st)<br>1/500 (2nd) | rabbit IgG |
| CD117<br>(c-KIT) | PE-Cy5 | BioLegend | 313210 | 1/50 | mouse IgG1 |
| CD123 | APC-Cy7 | Miltenyi | 130-115-267 | 1/50 | recombinant<br>human IgG1 |
| GPR56 | APC | Biolegend | 391906 | 1/100 | mouse IgG1 |
| CD34 | FITC | BD Biosciences | 555821 | 1/50 | mouse IgG1 |
| CD38 | PE-Cy7 | BD Biosciences | 560677 |  | mouse IgG1 |
| CD200 | BV711 | BioLegend | 329222 | 1/50 | mouse IgG1 |
| CLL-1 | BV605 | BD Biosciences | 742931 |  | mouse IgG2a |
| MRC2 | PE | BD Biosciences | 566817 | 1/100 | mouse IgG1 |
| **AML phenotyping (ImageStream)** |  |  |  |  |  |
| CD33 | APC | BD Biosciences | 551378 | 1/50 | mouse IgG1 |
| CD34 | APC | BD Pharmagen | 555824 | 1/50 | mouse IgG1 |
| Integrin-α7 | Unconjugated // gt-a-rabbit Alexa 546 | Thermo Fisher / invitrogen | PA5-49447 // A11010 | 1/150<br>1/500 | rabbit IgG |
| viability | BV510 | Thermo Fisher | L34966 | 1/2000 |  |
| DAPI |  | Fisher Scientific | D1306 | 1/2000 |  |

### ImageStream analysis

Extracellular antigens were stained as described for flow cytometry. Subsequently, cells were fixed and permeabilized using the Cytofix/Cytoperm kit (BD Biosciences, 554722) as recommended by the manufacturer. Fixed cells were blocked in Perm/Wash Buffer (BD, 554723), followed by intracellular staining of integrin α7 and DAPI (Table 1). Cells were acquired using the same laser settings on the Cytek Amnis ImageStream Mk II imaging flow cytometer (Cytek Biosciences). For gating strategies see **Supplemental Figure S6**.

### Gene expression analysis using quantitative real-time PCR

Total RNA was extracted using the RNeasy Mini kit (Qiagen, 74106) according to the manufacturer’s instructions. cDNA was synthesized from 600 ng total RNA using the High-capacity cDNA Reverse Transcription Kit (ThermoFisher, 4368814) according to the manufacturer’s instructions including 25U RNA Inhibitor (Roche/Merck, 3335399001) per reaction. cDNA was diluted 1:3 and qRT-PCR was conducted using a SYBR Green master mix (Roche, FSUSGMMRO) on a Light Cycler 480 (Roche). Gene expression was normalized to the expression of GAPDH and TBP.

### Western blotting

Protein lysates were prepared from 5x10^6^ cells using cell lysis buffer (Cell Signaling Technology, 9803S) supplemented with protease-phosphatase inhibitor (ThermoFisher, 78442) as recommended by the manufacturer. 20 µg denatured protein lysates were loaded on 10 % SDS-polyacrylamide gels and separated by SDS-PAGE.

Proteins were transferred to a nitrocellulose membrane (BioRad, 1705158) using the transblot Turbo Transfer System (BioRad, 1704150). The membranes were blocked for 1 h with 5 % non-fat dry milk (Cell Signaling Technology, 9999) in TBST. The membranes were probed with antibodies depicted in table 2. Chemiluminescent protein detection reagents were applied for GAPDH (Merck, GERPN2232) and laminin receptors (Thermo Fisher Scientific, 34094). Protein signals were acquired with a Fusion FX imaging system (Vilber).

**Table 2:** Antibodies for Western blotting.

| Marker | Species | Manufacturer | Order No. | Dilution | Size |
| --- | --- | --- | --- | --- | --- |
| **Primary** |  |  |  |  |  |
| BCAM | mouse | R&D biotechne | MAB1481 | 1/500 | 70-90 kDa |
| GAPDH | rabbit | Cell Signaling | 2118 | 1/2000 | 37 kDa |
| Integrin-α3 | rabbit | invitrogen | PA5-99407 | 1/500 | 117 kDa |
| Integrin-α6 | rabbit | invitrogen | 710209 | 1/1000 | 125, 150 kDa |
| Integrin-α7 | rabbit | Thermo Fisher | PA5-49447 | 1/2000 | 95, 125 kDa |
| **Secondary** |  |  |  |  |  |
| a-mouse HRP | horse | Cell Signaling Technology | 7076 | 1/3000 |  |
| a-rabbit HRP | goat | Cell Signaling Technology | 7074 | 1/3000 |  |

**Table 3:** Blocking antibodies.

| Marker | Manufacturer | Order No. | Dilution | Isotype |
| --- | --- | --- | --- | --- |
| anti-BCAM | R&D Systems | MAB1481 | 1/25 | mouse IgG2a |
| anti-integrin β1 | Merck | MABT821 | 1/25 | rat IgG2a |
| anti-integrin-α6 | BioLegend | 313602 | 1/25 | rat IgG2a |
| anti-integrin-α7 | Haas et al. 2017 (45) | | 2.5 μg | mouse IgG2a |
| **Isotype controls** |  |  |  |  |
|  | BD Biosciences | 562439 | 1/25 | mouse IgG2a |
|  | BioLegend | 400547 | 1/25 | rat IgG2a |

**Table 4:**
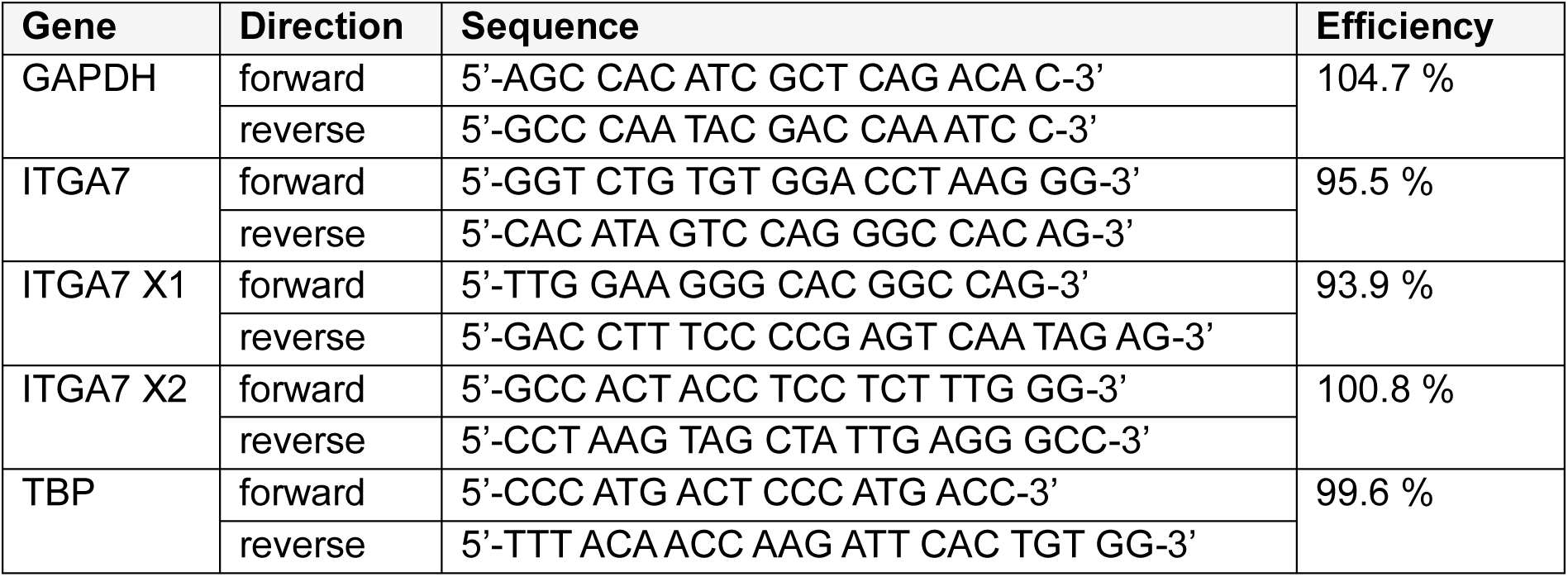
Primers for quantitative real-time PCR.

**Table 5:**
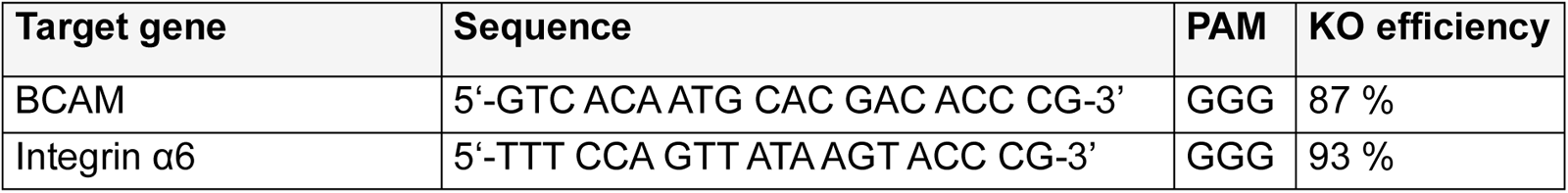
sgRNA sequences.

**Table 6:** primary AML samples.

| Patient | Sex | PB / BM | ELN 2017 | Integrin α3 [%] | Integrin α6 [%] | Integrin α7 [%] | BCAM [%] |
| --- | --- | --- | --- | --- | --- | --- | --- |
| 1 |  | PB |  | 0.31 | 21.9 | 46 |  |
| 2 | male | PB | adverse | 0.06 | 27.3 | 59.9 | 32.3 |
| 3 | male | PB | adverse | 0.08 | 73 | 37 | 80.3 |
| 4 | male | BM |  | 0.11 | 42.2 | 44 | 62.4 |
| 5 | female | PB | intermediate | 1.07 | 50.3 | 36.6 |  |
| 6 | female | PB | favorable | 0.13 | 66.1 | 66.1 | 4.53 |
| 7 | female | PB | favorable | 1.19 | 8.12 | 76.8 | 10.3 |
| 8 | female | PB |  | 0.24 | 2.03 | 3.02 | 9.63 |
| 9 |  | PB |  | 0.89 | 8.98 | 66.1 | 0.01 |
| 10 | female | PB | favorable |  | 2.44 | 0.65 | 7.93 |
| 11 | female | BM | favorable |  | 13.8 | 0.61 | 17.2 |
| 12 | female | BM | adverse |  |  |  |  |
| 13 | female | PB | favorable |  | 1.11 | 1.62 | 0.29 |
| 14 | female | BM | favorable |  | 0.81 | 1 | 2.24 |
| 15 | female | BM | intermediate |  |  |  |  |
| 16 |  | PB | intermediate | 1.78 | 37.5 | 31.5 | 49.1 |
| 17 | male | PB | favorable | 0.53 | 29.7 | 15.5 |  |
| 18 | female | PB | favorable | 0.04 | 22.6 | 5.91 |  |
| 19 | male | PB | favorable | 0.2 | 2.03 | 4.85 | 1.32 |
| 20 |  | PB | intermediate | 0.13 | 11.8 | 80.4 | 9.59 |
| 21 |  | PB |  | 0.08 | 33.8 | 64.8 | 31.9 |
| 22 | male | PB |  | 0.39 | 15.2 | 36.3 | 27.2 |
| 23 | male | PB |  | 0 | 2.44 | 0.32 | 4.93 |
| 24 |  | BM |  | 0.06 |  |  | 74.5 |
| 25 |  | PB |  | 0.08 | 0.09 | 31.9 |  |
| 26 |  | PB |  | 0.96 | 4.06 | 90.5 |  |
| 27 |  | PB |  | 0.26 | 25.1 | 0.16 |  |
| 28 | male | PB | intermediate |  |  |  | 0 |
| 29 | male | BM | intermediate | 4.51 | 11.2 | 0.16 | 0 |
| 30 | female | BM | favorable | 0.8 | 38.1 | 29.1 | 39.7 |
| 31 | female | BM | adverse | 3.64 | 43.5 | 16.1 | 3.89 |
| 32 | male | BM | adverse | 0.55 | 14.3 | 68.5 | 24.9 |
| 33 | male | PB | adverse | 3.67 | 70.4 | 72.8 | 46.5 |
| 34 | male | BM | adverse | 5.58 | 26.9 | 35.2 | 50.8 |
| 35 | female | PB | favorable | 0 | 0.55 | 3.52 | 8.79 |
| 36 | female | BM | favorable | 0.03 | 0.46 | 1.93 | 12.2 |
| 37 | female | PB | adverse | 0.14 | 63.5 | 86.8 | 59.1 |
| 38 | female | BM | adverse | 0.19 | 42.2 | 58.5 | 39.8 |
| 39 | male | BM | adverse | 2.32 | 44.6 | 17.9 | 52.7 |
| 40 | male | PB | favorable | 0.14 | 66.7 | 46.9 | 82.2 |
| 41 | male | BM | favorable | 1.51 | 37.4 | 30.5 | 37.6 |
| 42 | female | PB | intermediate | 0.23 | 64.6 | 17.4 | 43.3 |
| 43 | female | BM | intermediate | 0.17 | 81.9 | 8.5 | 60.9 |
| 44 | male | PB |  | 0.17 | 10.5 | 3.22 | 61.8 |
| 45 | female | PB | adverse | 0.08 | 1.07 | 1.39 | 8.53 |
| 46 | male | PB | favorable | 0.52 | 20.3 | 92.5 | 21.8 |
| 47 | male | PB | favorable | 0.61 | 14.1 | 8.3 | 0.27 |
| 48 | male | PB | adverse | 1.49 | 36.2 | 78.9 | 36.7 |
| 49 | female | PB | adverse | 3.88 | 10 | 2.94 | 30 |
| 50 | female | PB | favorable | 0.67 | 2.59 | 1.68 | 32.8 |
| 51 | male | PB | favorable |  |  |  | 7.36 |
| 52 | male | PB | adverse | 0.86 | 17.8 | 1.1 | 33.2 |
| 53 | male | PB | adverse |  |  |  | 1.07 |
| 54 | male | PB | intermediate | 0.28 | 1.25 | 1.46 | 3.1 |
| 55 | male | PB | favorable | 0.76 | 60.2 | 94.8 | 9.25 |
| 56 | female | PB | adverse | 6.67 | 68.8 | 67.9 | 67.5 |
| 57 | female | PB | adverse | 0.11 | 1.06 | 84.7 | 20.4 |
| 58 |  | PB |  | 0.08 | 74.5 |  |  |
| 59 |  |  |  |  |  |  | 0.92 |
| 60 |  |  |  |  |  |  | 2.58 |

### Colony-forming unit assays

Primary AML cells were sorted for laminin receptor expression. For integrin α6, α7 and BCAM, 80 000 sorted positive and negative cells were resuspended in 100 µL RPMI medium, transferred into 4 mL methylcellulose (Stem Cell Technologies, 4434) and vortexed vigorously. For integrin α3, the whole CD33+ cell fraction was used in comparison to CD33+ cells depleted for integrin α3. The cell suspension was incubated for 5 min at RT, before 20 000 cells were plated in triplicates into 35 mm petri dishes using a syringe and needle. Colonies were counted after 14 days.

### Transmigration assay

8 µm transwell membranes (Sarstedt, 83.3932.800) were coated with 0.05 % gelatin for 30 min at 37°C and washed with PBS before 50 000 HUVEC were seeded. HUVEC were cultured in endothelial cell medium (Promocell, C-22010) for 2 days to reach 100 % confluence. Patient samples were thawed and 0.4x10^6^ cells were seeded in 200 µL medium (RPMI, 2 % FBS, 1 % penicillin-streptomycin) onto the transwell. The bottom compartment was filled with 750 µL RPMI medium + 20 % FBS to create an FBS gradient or RPMI medium + 2 % FBS as control.

The transmigration assay was performed for 24 h at 37°C. Non-migrated cells were collected from the top compartment and migrated cells were collected from the 24 well plate. The transwell and the 24 well plate were rinsed with PBS to collect all remaining cells. For phenotypic analysis, cells were stained for flow cytometry. For quantitative analysis of cells previously sorted for integrin-α7 expression, cells were washed, resuspended in an equal volume of FACS buffer (PBS, 10 % FBS, 0.5 M EDTA) and acquired for a defined time at the same flow rate on a BD LSR Fortessa.

### Adhesion assay

For laminin coating, 2 µL laminin-111, laminin-211, laminin-332, laminin-411 and laminin-511 (BioLamina, LN111-02, LN211-02, LN332-02, LN411-02, LN511-0202, 30 µg/mL diluted with PBS + Ca2+/Mg2+) were applied in separate drops and in duplicates on 35 mm dishes (Greiner, P5112-740EA) and incubated for 45 min at RT. The dishes were carefully rinsed with PBS + Ca2+/Mg2+ and blocked with 5 % BSA in PBS + Ca2+/Mg2+ for 1 h at 37°C to prevent unspecific binding to plastic. 7x10^5^ cells were seeded in adhesion medium (RPMI, 10 % FBS, Ca2+, Mg2+, Mn2+) and incubated for 3 h at 37°C to allow cell adhesion. The cell suspension was aspirated and the dishes were very carefully rinsed 3x with PBS to remove non-adherent cells. Image acquisition was performed using a Zeiss Primovert microscope, brightness and contrast were adjusted with ImageJ (Version 1.53).

### Proliferation assay

A 96 well plate was coated with 50 µL laminin solution per well (BioLamina, LN211-02, LN511-0202, 10 µg/mL diluted with PBS + Ca2+/Mg2+) for 16 h at 4°C. The wells were washed 2x with PBS + Ca2+/Mg2+ before 20 000 SKM-1 AML cells were seeded per well. To determine the proliferation rate, cell counting kit 8 (Merck, 96992) was applied for 2.5 h as recommended by the manufacturer.

### RNA sequencing

Total RNA was extracted from CD33+ ITGA7-sorted primary AML cells (> 4x10^5^ cells/sample) using the RNeasy Mini kit (Qiagen, 74106) according to the manufacturer’s instructions. Extracted RNA was sent to Novogene Company Limited (UK) for library preparation and sequencing. Paired-end 150 bp sequencing was performed on an Illumina Novaseq 6000 with generation of 6 Gb per sample. Sequencing reads were mapped to the human reference transcriptome (hg37) and read counts per gene were quantified with the nf-core/rnaseq analysis pipeline (version 3.10) (25). Subsequently, gene counts were used for paired differential gene expression analysis with iDEP96 (26).

### Gene Set Enrichment Analysis (GSEA)

GSEA was performed from differentially expressed genes ranked according to logFC using WebGestalt (27). Only signatures with FDR < 0.05 are shown.

### Heatmaps

Heatmaps were generated from normalized gene counts (tpm) using Heatmapper (28). Average linkage was chosen as clustering method and Spearman Rank Correlation as distance method.

### Kaplan-Meier Survival Plots

KM-plotter was used to compare survival between patients with high and low gene expression in untreated samples from the datasets GSE1159 and GSE6891. The cutoff to separate samples into high and low expression was determined as described by Győrffy et al. (29).

### Statistics

Statistical analyses were performed using the GraphPad Prism software 9. The difference between two data sets was assessed using the non-parametric Mann-Whitney test or unpaired test with Welch’ correction depending on the sample distribution. For multiple comparisons, one-way Anova or two-way Anova with correction for multiple comparisons were performed. The Spearman’s correlation coefficient was used to calculate correlation between two variables. A p value below 0.05 indicates significance. *p<0.05; **p<0.01; ***p<0.001; ****p<0.0001.

## RESULTS

### Integrin α6 and α7 are overexpressed in AML

To characterize the laminin receptor profile in AML, we first compared publicly available transcriptome data from AML patients (TCGA) and healthy donors (GTEx) (30). We observed a significant overexpression of integrin α6 and α7 in AML compared to healthy BM while no significant difference was observed for BCAM and integrin α3 (**Figure 1A**). To confirm laminin receptor expression on the protein level, we performed multicolor flow cytometry on primary samples from healthy donors (mobilized CD34+CD38-HSPC, n=5), AML patients at diagnosis (n=60) and AML cell lines (n=7). Overall, we observed a very heterogeneous expression of surface antigens among CD33+ bulk AML samples (**Figure 1B**). Integrin α3 displayed the lowest expression on bulk AML cells, which was comparable to the surface expression on CD34+ CD38-HSPC. In contrast, higher mean expression was found for integrin α6, α7 and BCAM with integrin α6 and α7 again showing a trend towards upregulation on AML cells compared to HSPC (**Figure 1B**). In general, the expression patterns furthermore differed among individual AML patient samples, with some patients showing two distinct populations and others rather a continuum of laminin receptor expression (**Figure 1C**). While integrin α7 staining often allowed to distinguish a positive and negative population, integrin α6 and BCAM staining in most cases exhibited a continuum from negative to positive expression. AML cell lines also displayed various expression profiles. While most cell lines expressed a combination of different laminin receptors, the cell line MOLM-13 was negative for all analyzed laminin receptors **(Supplemental Figure S1A)**. However, the absence of surface expression did not necessarily correspond to the absence of intracellular protein. Particularly, MOLM-13 and Kasumi-1 showed a discrepancy regarding integrin α7, which only showed intracellular localization in these cell lines **(Supplemental Figure S1B-C)**. Interestingly, integrin α7 was the only laminin receptor showing differential expression between AML samples from PB and BM with higher expression on PB cells **(Supplemental Figure S1D)**.

**Figure 1.**
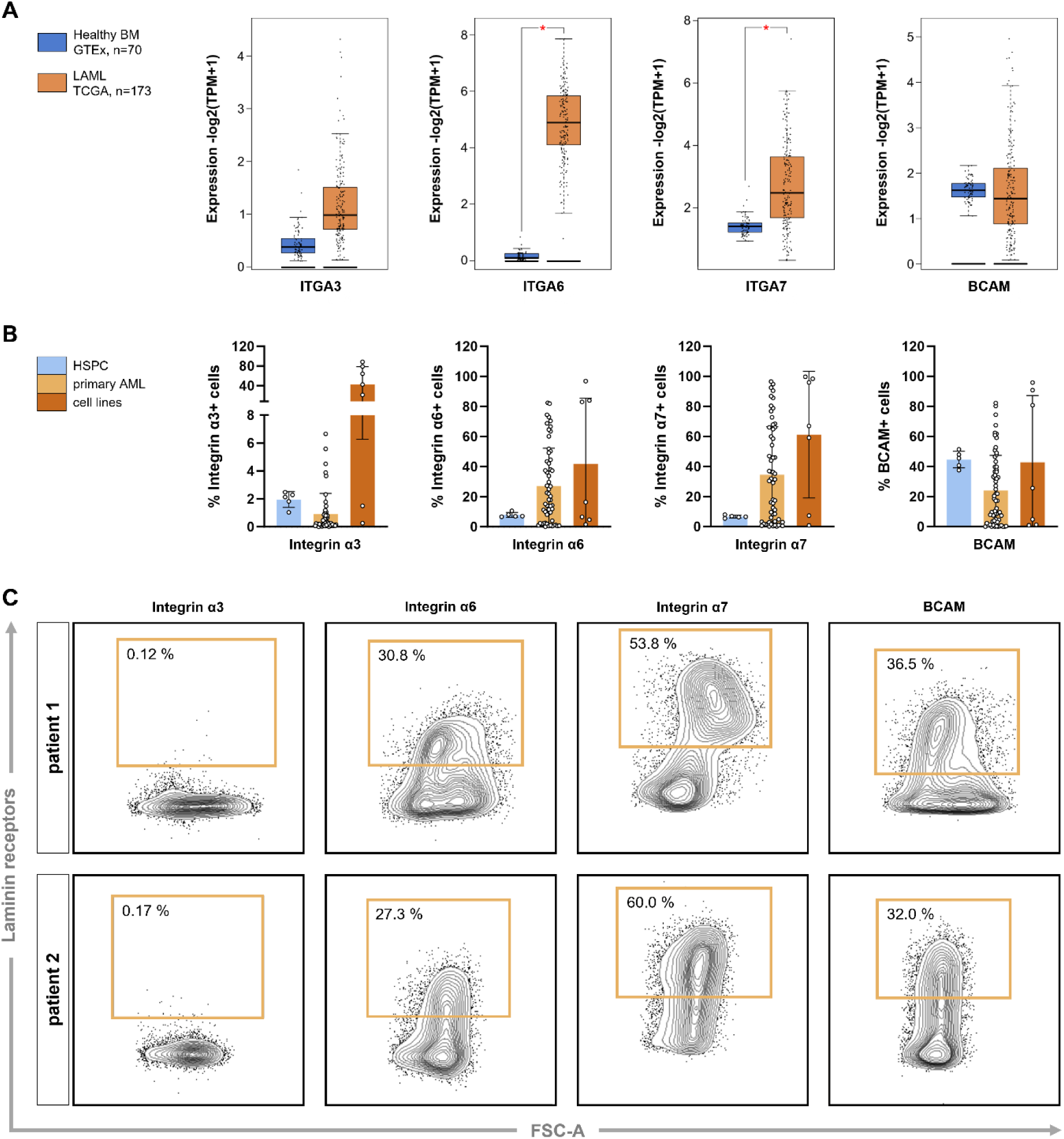
Characterization of laminin receptor expression reveals an overexpression of integrin α6 and α7 in AML. **(A)** Comparison of gene expression between healthy BM samples (GTEx, n=70) and AML samples (TCGA, n=173) using Gepia (30). ITGA6 and ITGA7 are overexpressed in AML. **(B)** Flow cytometry analysis of laminin receptor expression on healthy mobilized CD34+ CD38-HSPC (n=5), primary CD33+ AML cells (n=60) and AML cell lines (n=7) shows an overexpression of integrin α6 and α7 in AML, depicted as mean +/-SD. **(C)** Exemplary flow cytometry staining of two selected patients showing laminin receptor surface expression on primary AML cells.

### Integrin α7 is associated with differentiation and its absence enriches for LSC

The interaction of LSC with the BM niche is particularly critical, as it is thought to mediate the survival of this disease-driving subpopulation (31). To identify potential differences in how LSC and non-LSC interact with the BM, we sought to investigate the distribution of laminin receptors on LSC versus non-LSC populations of primary AML samples by using co-staining with established markers distinguishing these populations, such as CD34, NKG2DL, cKit and GPR56 (32). Our data indicate that only integrin α3 is enriched on LSC (CD34+), albeit at a low overall expression level. In contrast, integrin α7 was specifically enriched on non-LSC (CD34-) (**Figure 2A**). Integrin α7 also exhibited enhanced expression on CD34-non-expressing AML (<1 % CD34+ cells) compared to CD34-expressing AML (>5 % CD34+ cells), whereas the opposite was found for integrin α6 and BCAM **(Supplemental Figure S1E)**.

**Figure 2.**
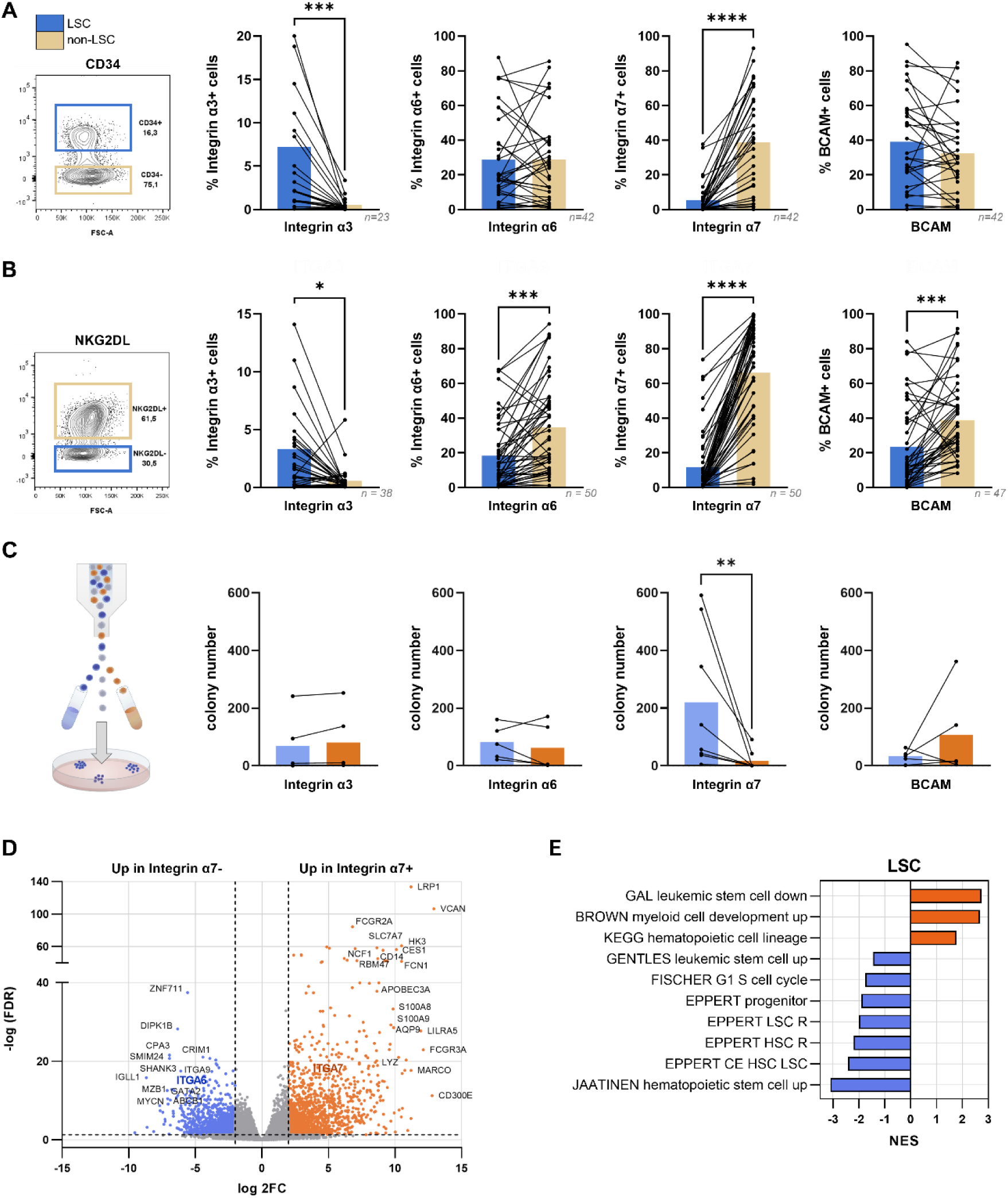
Integrin α7 is enriched in non-LSC and associated with myeloid differentiation. Co-expression of laminin receptors with LSC markers shows enrichment for integrin α3 in the LSC-compartment and enrichment for integrin α7 in the non-LSC compartment. **(A)** CD34 expression defines the LSC compartment. **(B)** The absence of NKG2DL defines the LSC compartment. **(C)** Colony-forming unit (CFU) assays from primary AML cells sorted for laminin receptor expression reveal higher clonogenic activity of integrin α7-cells compared to their integrin α7+ counterpart. In contrast to the other laminin receptors that were sorted for positive and negative populations, integrin α3 was depleted from CD33+ cells, which served as control. **(D)** Four primary AML samples were sorted for integrin α7 surface expression and RNA-sequencing was performed for both cell fractions. Vulcano plot of differentially expressed genes (logFC > 2, FDR < 0.05). **(E)** GSEA of RNA-sequencing data for LSC-related genes. Stemness-related signatures are enriched in integrin α7-cells whereas differentiation signatures are enriched in integrin α7+ cells.

We additionally used the expression of NKG2DL to define LSC (NKG2DL-) and non-LSC (NKG2DL+) populations. This approach allows to distinguish LSC and non-LSC populations regardless of CD34 expression status (33). Here, we similarly found that integrin α3 was enriched on LSC, whereas the integrins α6, α7, and BCAM were enriched on non-LSC (**Figure 2B**). Consistently, integrin α7 was the only laminin receptor that was also negatively correlated with other LSC markers such as c-Kit or GPR56 **(Supplemental Figure S2A)**.

To determine whether the differential laminin receptor expression marks functionally distinct populations, we performed colony-forming unit (CFU) assays of primary AML samples sorted according to their laminin receptor expression. Only integrin α7 was able to distinguish populations with different clonogenic potential. As expected, integrin α7-cells had a significantly higher clonogenic potential than integrin α7+ cells, indicating a higher frequency of non-LSCs in the latter subpopulation (**Figure 2C**).

#### Transcriptomic analysis reveals differentiation signatures in integrin α7+ cells

Next, we investigated gene expression differences between integrin α7+ and integrin α7-subpopulations from 4 primary AML samples. Indeed, distinction based on integrin α7 surface expression also allowed to distinguish transcriptionally different populations **(Supplemental Figure S2B)**. Transcriptomic analysis identified 2 506 differentially expressed genes (DEG, |log2FC|>2, FDR<0.05) with 1 195 genes upregulated in integrin α7-cells and 1 311 genes upregulated in integrin α7+ cells (**Figure 2D**). Gene set enrichment analysis (GSEA) confirmed that the absence of integrin α7 was associated with stemness signatures, whereas integrin α7 was associated with differentiation signatures (**Figure 2E and Supplemental Figure S2C)**. Furthermore, integrin α7+ cells showed enrichment of biological processes involved in immune response and inflammation that also occur during differentiation **(Supplemental Figure S2D)**. In line with this observation, integrin α7 surface expression was associated with a differentiation signature in healthy cord blood samples. In contrast to healthy HSPC with mainly intracellular integrin α7 localization, healthy PBMC and AML cells showed the most pronounced integrin α7 staining on the cell surface **(Supplemental Figure S3A-C)**. This makes integrin α7 an interesting target to distinguish HSPC from AML cells. In summary, our results show that in line with a higher expression on more differentiated healthy cord blood cells integrin α7 reliably identifies non-LSC in AML.

### Integrin α7 marks a cell population with migratory capacity

To gain insight into the role of integrin α7 in AML, we aimed to identify molecular signatures that are not only linked to differentiation but allow a better understanding of integrin α7+ cell function. To this end, we generated an integrin α7-specific transcriptomic signature by comparing our RNA-sequencing data with our previously generated transcriptomic dataset (GSE127959) that distinguishes LSC and non-LSC populations (|log2FC|>1)(33). GSEA was performed using the genes that displayed differential expression solely in our integrin α7 dataset (**Figure 3A**). Interestingly, GSEA indicated a potential involvement of integrin α7+ cells in processes such as cell migration, angiogenesis and endothelial-to-mesenchymal transition (EMT) (**Figure 3B, Supplemental Figure S4)**. To determine whether integrin α7 contributes to cell migration, we employed transmigration assays across a confluent endothelial cell layer. When using primary AML cells sorted for integrin α7 expression, those assays revealed a higher migratory capacity of integrin α7+ cells compared to their integrin α7-counterparts (**Figure 3C**). Similarly, phenotyping of unsorted primary AML cells after transmigration assays revealed a higher expression of integrin α7 on migrated cells compared to non-migrated cells (**Figure 3D**). When taking a closer look at our transcriptomic dataset, we found 79 DEGs in the integrin α7+ compartment that are also integrin-related genes (**Figure 3E**). Strikingly, many of these genes such as versican (*VCAN*), thrombospondin-1 (*THBS1*) or syndecan-2 (*SDC2*) are also associated with cell migration (34,35) further supporting our hypothesis that integrin α7 identifies a migratory leukemic cellular subpopulation (**Figure 3F**).

**Figure 3.**
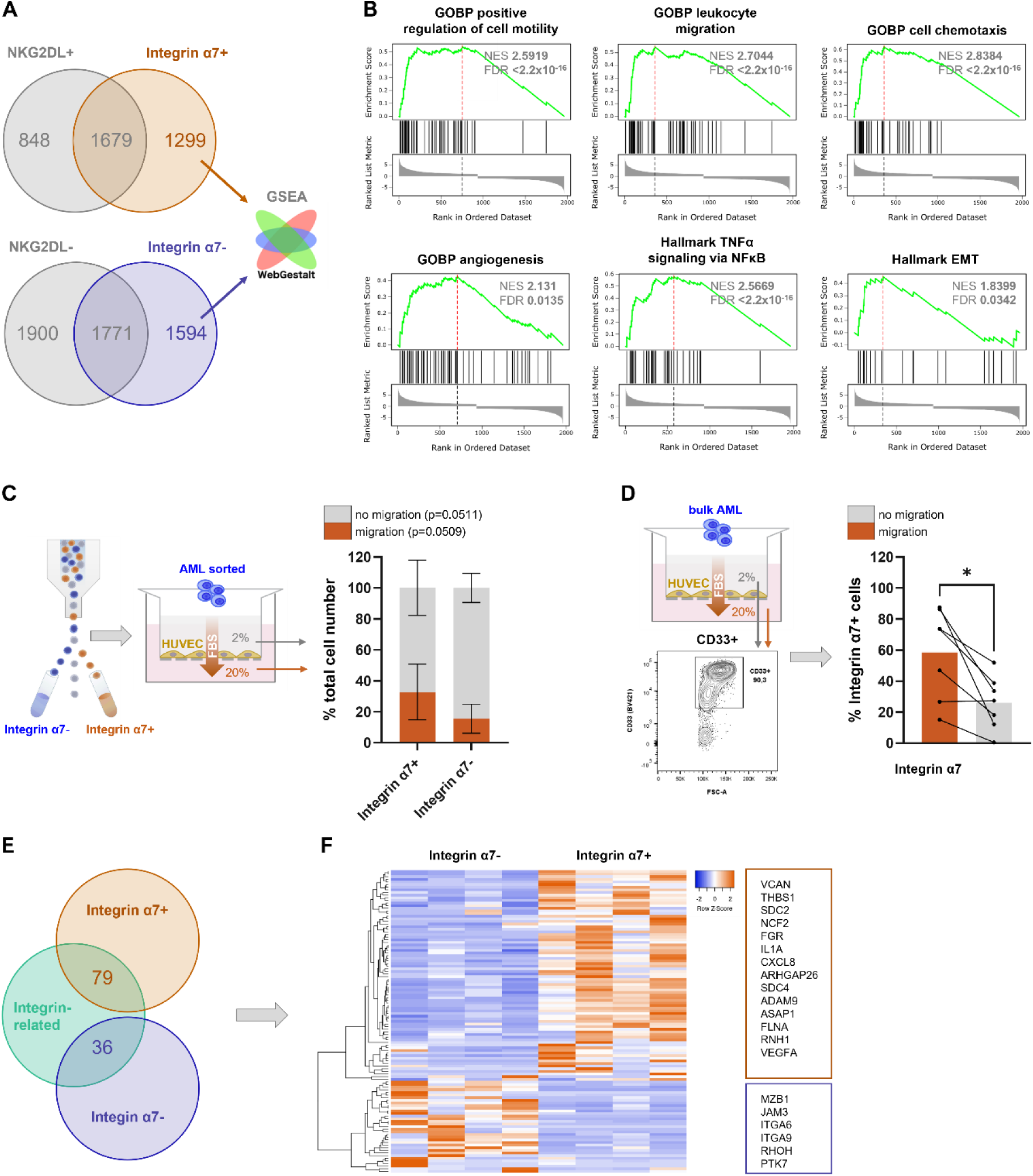
Integrin α7 defines a cell population with migratory capacity. **(A)** Differentially expressed genes (logFC > 1) were compared to an LSC-dataset based on NKG2DL expression (logFC > 1). The genes unique to the integrin α7 dataset were used for GSEA. **(B)** GSEA of genes unique to the integrin α7 dataset reveals enrichment of molecular signatures related to cell migration, angiogenesis and EMT. **(C)** Transmigration assays of primary AML cells were performed across a confluent layer of endothelial cells (HUVEC). Primary AML cells were sorted for integrin α7 surface expression and both cell fractions were seeded for transmigration assays. Migrated and non-migrated cells were counted after 24 h. **(D)** Bulk AML cells were seeded for transmigration assays and both, migrated and non-migrated cell fractions were analyzed via flow cytometry after 24 h. Phenotyping shows higher integrin α7 expression on migrated cells compared to non-migrated cells. **(E)** Genes were extracted from integrin-related molecular signatures from the Molecular Signature Database (Human MSigDB v2023.2) (52,53) and overlapped with differentially expressed genes (logFC > 2, FDR < 0.05). **(F)** Heatmap of integrin-related genes in integrin α7- and integrin α7+ cells.

### AML cells strongly adhere to laminin-511 and weakly to laminin-332

Since the involvement of integrins in cell migration requires cell adhesion, we next investigated whether integrin α7 also contributes to laminin adhesion. Initially, we performed adhesion assays to the laminin isoforms LM-511, -411 and -332 described to be secreted in the BM (36–38). Overall, we observed strong adhesion of primary AML cells and AML cell lines to laminin-511, weak adhesion to laminin-332 and no adhesion to laminin-411 (**Figure 4A and Supplemental Figure S5A)**. However, laminin-specific adhesion was not displayed in all primary AML samples **(Supplemental Figure S5B)**. Of note, laminin-511 represents a ligand described as being bound by all the analyzed laminin receptors (**Figure 4B**) (15,39). Flow cytometric analyses of adhesive and non-adhesive cells indicated a tendency towards greater integrin α6 and α3 expression and decreased NKG2DL expression in laminin-511 adherent cells (**Figure 4C**). Laminin-211 has previously been shown to increase the proliferation of integrin α7-positive cell lines (24). Therefore, we investigated whether integrin α7 mediates adhesion to laminins other than those described in human BM. However, our adhesion assays revealed that neither primary AML cells nor AML cell lines exhibited adhesion to laminin-211 or laminin-111 (**Figure 4D**). Furthermore, we only observed a slightly increased proliferation of SKM-1 cells in the presence of laminin-211 (**Figure 4E**).

**Figure 4.**
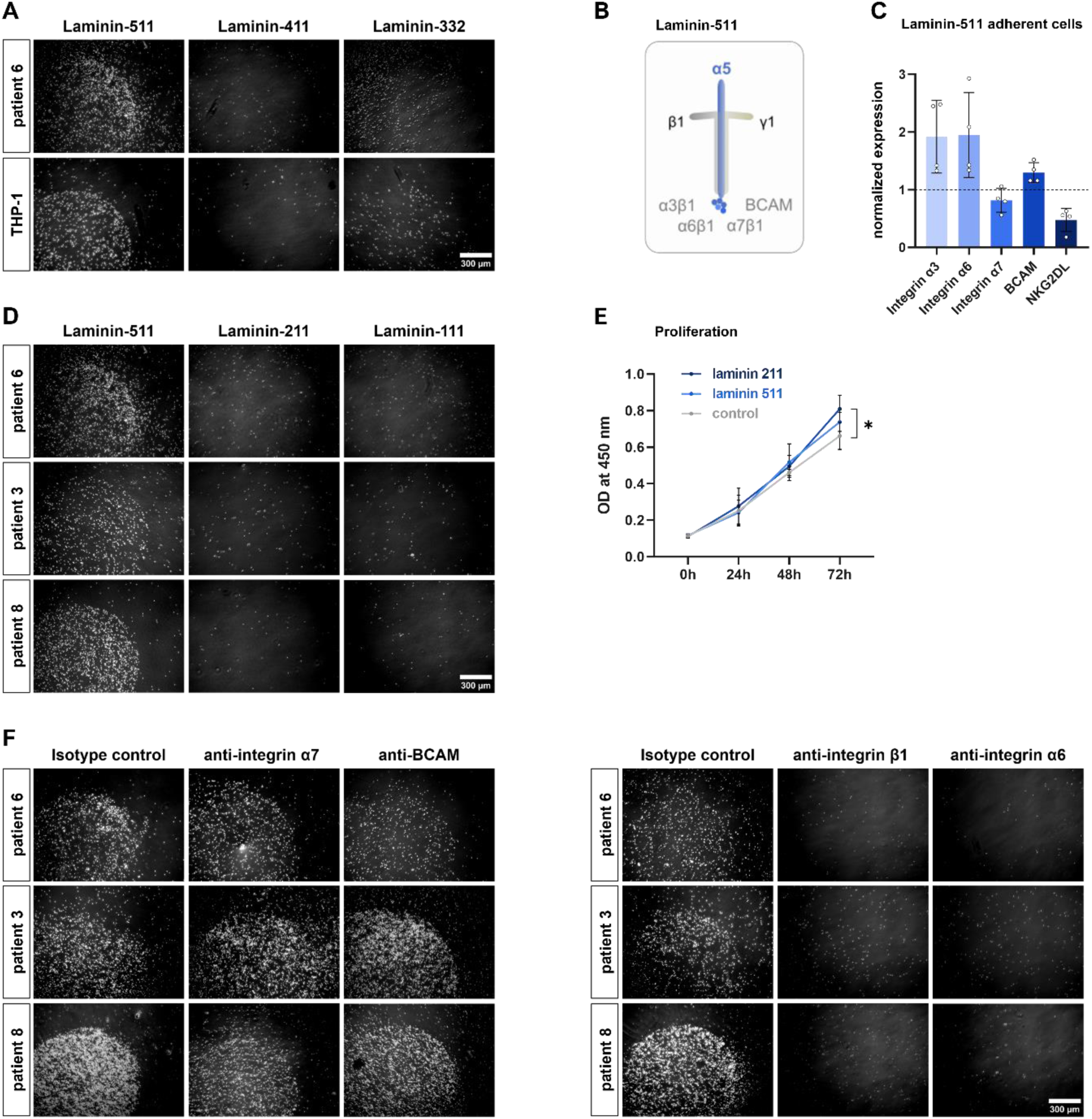
Blocking of integrin α6 but not integrin α7 abrogates AML cell adhesion to laminin-511. **(A)** Representative images showing adhesion assays with primary AML cells and the AML cell line THP-1 on different laminin isoform coatings. Plastic dishes were coated with single laminin isoform droplets and unspecific cell adhesion to plastic was blocked with BSA. Laminin specific adhesion is seen in the circular droplet region. Brightfield images were acquired with a microscope (Zeiss, Primovert). **(B)** Schematic drawing of laminin-511 and its binding specificities. All four laminin receptors have been described to bind to the C-terminal end of laminin-511 (15,39) **(C)** Phenotyping of adherent and non-adherent cell fractions, marker expression on adherent cells was normalized to the expression on non-adherent cells within each sample. Adherent cells show a higher expression of integrin α3 and α6 and a reduced expression of NKG2DL. **(D)** Primary AML cells adhere to laminin-511 but not to laminin-211 or laminin-111. **(E)** Proliferation assays with the SKM-1 AML cell line on different laminin coatings or without coating (control). Laminin-211 coating slightly increases cell proliferation (n=3). **(F)** Blocking antibodies against integrin α6 and integrin β1 reduce adhesion of primary AML cells to laminin-511 whereas blocking antibodies against integrin α7 and BCAM have no effect on cell adhesion.

### Integrin α6 mediates laminin adhesion in AML

While blocking antibodies against integrin α7 and BCAM did not affect adhesion to LM-511, the adhesive interaction was abrogated by blocking antibodies against integrin α6 and integrin β1 (**Figure 4F**). Similarly, deleting the integrin α6 chain in Kasumi-1 cells reduced adhesion to LM-511 **(Supplemental Figure S5C)**. Together, our data indicate that integrin α6 and integrin α7 distinguish two functionally separate populations in AML, the former with superior adhesive capacity, and the latter with greater migratory potential. Therefore, the question arises whether the integrins α6 and α7 are co-expressed in AML. While we observed a non-LSC population expressing both markers, LSC showed practically no co-expression, and the integrin α6+ integrin α7-compartment was significantly higher compared to the double positive compartment **(Supplemental Figure S5D)**.

### Laminin receptor expression is associated with poor prognosis

Since the interaction with the BM niche and the expression of receptors for such interactions can foster leukemia development (40), we evaluated laminin receptor expression in different risk groups classified according to the ELN2017 risk stratification (41). Interestingly, we observed a trend towards higher expression of integrin α6, α7, and BCAM in the adverse risk group as opposed to favorable samples (**Figure 5A**). Notably, high expression of all laminin receptors at diagnosis correlated with shorter survival in the GSE1159 and GSE6891 datasets (**Figure 5B**).

**Figure 5.**
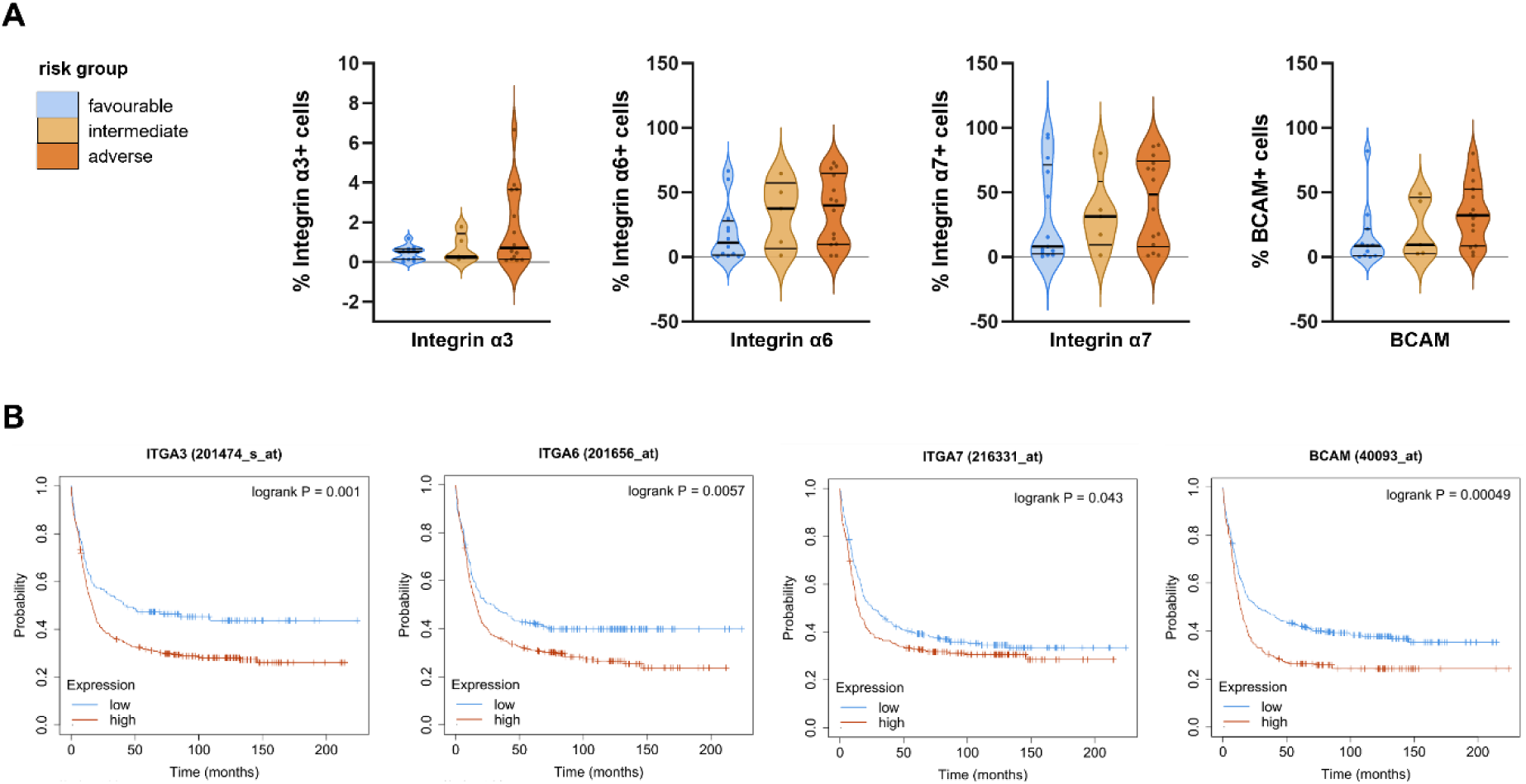
Laminin receptor expression is associated with poor prognosis in AML. **(A)** Patient samples were categorized according to the ELN2017 risk stratification and laminin receptor expression was analyzed via flow cytometry. Integrin α6, α7 and BCAM show a trend towards higher marker expression in the adverse risk group compared to the favorable risk group. **(B)** Kaplan-Meier plots showing differences in survival between untreated AML samples with high or low laminin receptor expression (mRNA level). The plots were generated from the datasets GSE1159 and GSE6891 using KM-plotter (29).

## DISCUSSION

It is widely acknowledged that interactions with the BM niche are essential for cell survival, self-renewal or drug resistance of leukemic cells, which therefore express a broad spectrum of surface molecules and receptors to engage with niche cells and ECM proteins, such as laminins (5,6,42,43). While laminin receptors have been associated with stemness in healthy hematopoiesis (18–20), their expression and function in AML remain understudied. Here, we provide a comprehensive characterization of the laminin receptors integrin α3β1, α6β1, α7β1 and BCAM in AML. Overall, we noted a heterogeneous laminin receptor expression on primary AML cells, with integrin α6 and α7 displaying the most abundant cell surface presentation and overexpression on mRNA level in comparison to healthy HSPC.

Previous studies have shown that solid tumor cancer stem cells express the integrins α6 and α7 (44–46). Furthermore, integrin α6-mediated laminin adhesion has been reported to promote quiescence in EVI1^high^ AML (22). Thus, we aimed to determine whether laminin receptors are also differentially expressed by stem and non-stem cell populations in AML. Using a flow cytometry approach, we noted that integrin α3 was enriched on LSC, while integrin α7 was absent on LSC. In addition, integrin α7 surface expression allowed to distinguish between LSC and non-LSC populations on both a transcriptional and functional level. It is important to note that this also applies to AML samples without CD34 expression. In such cases, the distinction between LSC and non-LSC compartment is more difficult due to the limited number of surface markers available to identify either subset. Interestingly, a discrepancy between total integrin α7 protein expression and its cell surface localization was noted in the AML cell lines MOLM-13 and Kasumi-1, as well as in healthy HSPC. Here, we only observed intracellular integrin α7 signals but no surface expression of the receptor. This indicates that protein localization may be differentially regulated in AML cells compared to HSPC. The high expression of integrin α7 on AML cells, combined with its absence on HSPC, makes it a potential target molecule in AML.

Regardless of whether the laminin receptors were found to be enriched in the LSC or the non-LSC compartments, analysis of online transcriptomic datasets indicated that the elevated expression of all four laminin receptors was associated with shorter overall survival. Of note, laminin receptor expression has previously been linked with prognosis. In particular, ITGA6 expression has been described to be higher in relapse samples compared to remission samples in EVI1^high^ AML, and high expression of ITGA7 has been associated with poor prognosis in AML (22,23).

Noteworthy, overexpression of integrin α7 has been reported in patients with myeloid sarcoma, an extramedullary AML manifestation, which has also been described as an independent risk factor in AML (24). The development of extramedullary disease requires the extravasation of AML cells from the BM and their migration to distant tissue sites. Remarkably, our analyses revealed higher integrin α7 surface expression on PB cells as opposed to BM cells, and our RNA-sequencing data furthermore indicated a potential involvement of integrin α7 in cell migration. Among the molecular signatures enriched in integrin α7+ cells we found processes such as cell migration, angiogenesis and EMT. In solid tumors, EMT is described as a cancer hallmark enabling tumor cells to migrate and metastasize (47). Although leukemia represents non-epithelial tumors, EMT-like expression patterns have been associated with poor prognosis in AML (48,49).

Functional studies revealed a higher migratory capacity of integrin α7+ cells compared to their integrin α7-counterpart. However, it is likely that additional surface proteins contribute to cell migration. Our RNA-sequencing data indicate that integrin α7+ cells express various genes with a potential involvement in cell migration. Notably, VCAN and THBS1 have been previously reported to promote leukemia cell migration (34,35). Therefore, it cannot be asserted that integrin α7 is directly involved in cell migration or simply marks a migratory cell population.

ECM proteins have been reported to impact cell proliferation directly and indirectly. ECM-integrin interactions have been shown to influence cell cycle progression, and laminin networks contain growth factors that act alone or in synergy with laminin-integrin interactions (10,50). Consistently, previous studies have shown a proliferative advantage of integrin α7+ cell lines in the presence of laminin-211 (24). We confirmed a mild increase in proliferation on laminin-211 coatings using the AML cell line SKM-1. However, AML cells did not adhere to laminin-211, despite the expression of the ITGA7 isoform α7X2, which shows a preferred binding to laminin-211 (15) **(Supplemental Figure S5E)**. While integrin α7 did not contribute to laminin adhesion in general, integrin α6 mediated strong adhesion to laminin-511, one of the predominant laminin isoforms in human BM.

Altogether, we suggest that integrin α7 surface expression defines a more mature cell population with migratory capacity and the potential to extravasate from the BM. In contrast, integrin α6 contributes to laminin adhesion and therefore presumably to AML cell retention to the BM niche. Laminin receptors may balance cell migration and adhesion, and further studies are needed to understand how different niches affect this balance. Interfering with ECM interaction and targeting the migratory cell population may have therapeutic potential in AML.

In particular targeting integrin α7 may be an interesting strategy regarding challenging cases with extramedullary AML (51).

## Data availability

Gene counts from RNA sequencing can be provided upon request.

## Acknowledgements

The authors thank the staff of the FACS core facility FCF Berg (University Hospital Tübingen) for cell sorting and advice. We thank Tobias Haas and Maria De Ruggero (Institute of General Pathology, Università Cattolica del Sacro Cuore, Rome, Italy) for kindly sharing the integrin α7 blocking antibody. We thank Jan Pauluschke-Fröhlich and Harald Abele (Frauenklinik Tübingen) for providing cord blood samples. We thank all patients for donating blood and BM. This project has received funding from the European Research Council (ERC) under the European Union’s Horizon 2020 research and innovation program (grant agreement n° 866548, HemStem, awarded to CL), the German Cancer Consortium (DKTK) Joint Funding Program (RiskyAML), the Deutsche Forschungsgemeinschaft (DFG, German Research Foundation) – 467578951, and the German Cancer Aid (NK fit against AML), and the DFG (91b grant for the BD Lyric). Supported by the BMBF-funded de.NBI Cloud within the German Network for Bioinformatics Infrastructure.

## Author contributions

EG and GK conceived the project and contributed to experimental design. EG performed experiments, analyzed and interpreted data. MA contributed to experimental work, data analysis and interpretation. MK analyzed RNA-sequencing raw data. JW provided information on the ELN2017 risk stratification of patient samples. EG prepared the manuscript with input from all authors. CL supervised the study.

## Abbreviations

A: ampere
ACK: ammonium–chloride–potassium
AML: acute myeloid leukemia
BCAM: basal cell adhesion molecule
BM: bone marrow
BSA: bovine serum albumin
Cas9: CRISPR associated protein 9
CB: cord blood
cDNA: complementary DNA
CFU: colony-forming unit
cm2: square centimeters
DAPI: 4′,6-diamidin-2-phenylindol
ddH2O: double-distilled water
DEG: differentially expressed genes
DMSO: dimethyl sulfoxide
ECM: extracellular matrix
EDTA: ethylene diamine tetraacetic acid
ELN: European leukemiaNet
EMT: epithelial-to mesenchymal transition
FACS: fluorescence-activated cell sorting
FBS: fetal bovine serum
FC: fold change
FDR: false discovery rate
FLT3-L: fms-related receptor tyrosine kinase 3 ligand
g: gram
GAPDH: glyceraldehyde 3-phosphate dehydrogenase
Gb: giga base
G-CSF: granulocyte colony-stimulating factor
GTEx: genotype-tissue expression
GSEA: gene set enrichment analysis
h: hours
HKG: housekeeping gene
HSPC: hematopoietic stem and progenitor cell
HUVEC: human umbilical vein endothelial cells
IL-3: interleukin-3
IL-6: interleukin-6
ITGA3: gene encoding integrin α3
ITGA6: gene encoding integrin α6
ITGA7: gene encoding integrin α7
KO: knockout
l: liter
LM: laminin
LSC: leukemic stem cell
µg: microgram
µL: microliter
mg: milligram
min: minutes
mL: milliliter
mM: millimolar
MNC: mononuclear cell
NFκB: nuclear factor kappa-light-chain-enhancer of activated B cells
NKG2DL: natural killer group 2D ligand
nm: nanometer
nmol: nanomol
OD: optical density
PB: peripheral blood
PBMC: peripheral blood mononuclear cells
PBS: phosphate-buffered saline
PCR: polymerase chain reaction
Pmol: picomol
qRT-PCR: quantitative real-time polymerase chain reaction
RNP: ribonucleoprotein
RT: room temperature
SCF: stem cell factor
SDC2: syndecan-2
SDS: sodium dodecyl sulfate
SDS-PAGE: sodium dodecyl sulfate polyacrylamide gel electrophoresis
sgRNA: single guide RNA
SR-1: StemRegenin 1
TBP: TATA-binding protein
TBST: Tris-buffered saline with Tween20
TCGA: the cancer genome atlas
THBS1: thrombospondin 1
TNFα: tumor necrosis factor alpha
tpm: transcripts per million
TPO: thrombopoietin
UM729: pyrimido-indole derivate
V: volt
VCAN: versican

## Supplementary

**Supplemental Figure S1.**
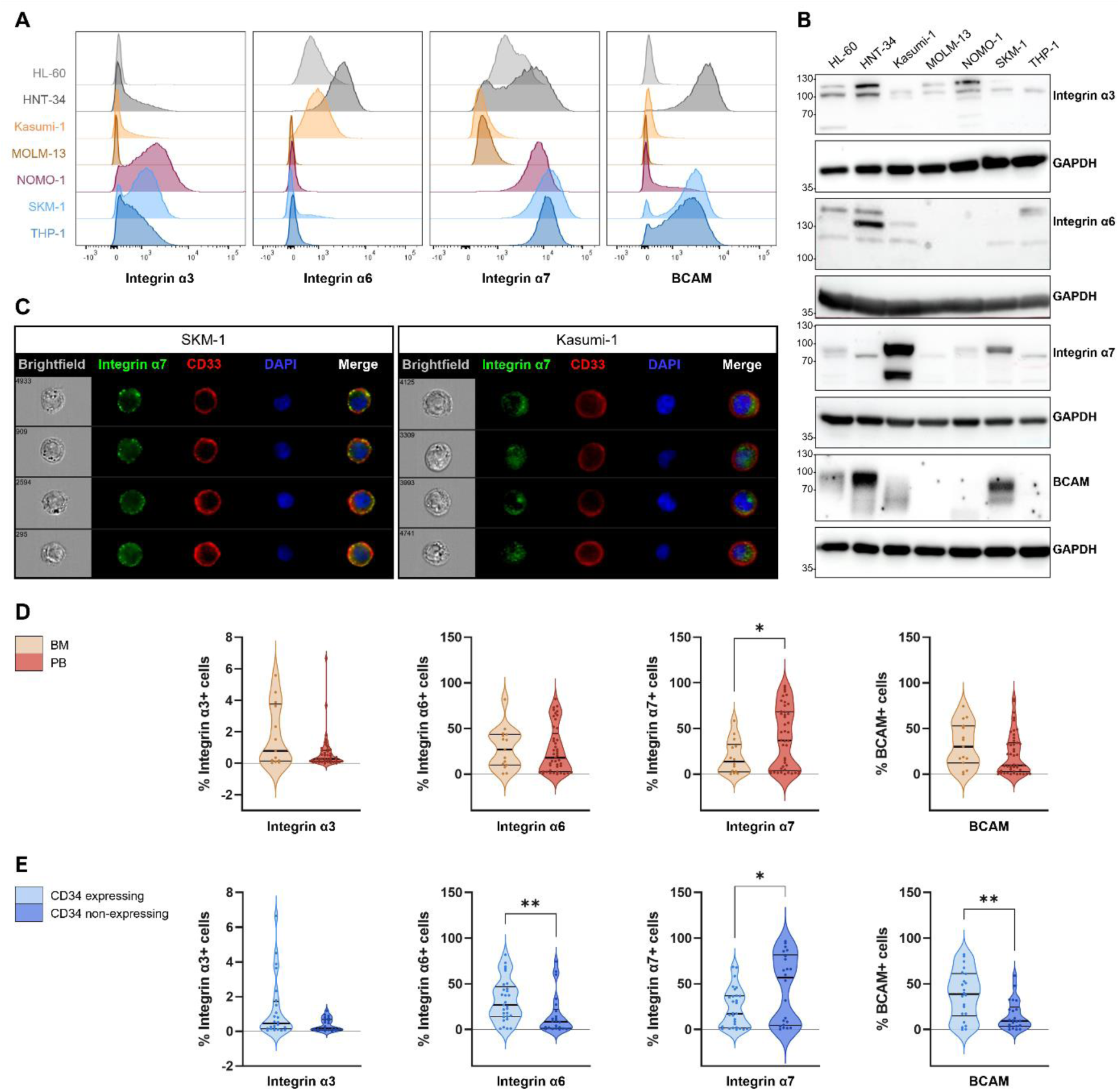
Laminin receptor expression in AML cell lines and primary AML cells. **(A)** Flow cytometric analysis shows heterogeneous laminin receptor surface expression between AML cell lines. HNT-34, SKM-1 and THP-1 display the expression of the most laminin receptors as opposed to MOLM-13, which is negative for all laminin receptors analyzed. **(B)** Western blotting of laminin receptors confirms the expression patterns of integrin α3, α6 and BCAM as seen by flow cytometry. Expression of integrin α7 is very pronounced for Kasumi-1 in Western blotting, but not measurable via flow cytometry. **(C)** ImageStream analysis of intracellular and extracellular integrin α7 shows expression on the cell surface in SKM-1, but only intracellular expression in Kasumi-1. **(D)** Comparison between laminin receptor expression on BM and PB primary AML cells reveals higher integrin α7 expression on PB cells. **(E)** Comparison between laminin receptor expression on AML samples positive for CD34 (> 5 % CD34+ cells) and negative for CD34 (< 1 % CD34+ cells) shows higher surface expression of integrin α7 on CD34-AML samples, whereas integrin α6 and BCAM are higher expressed in CD34+ AML.

**Supplemental Figure S2.**
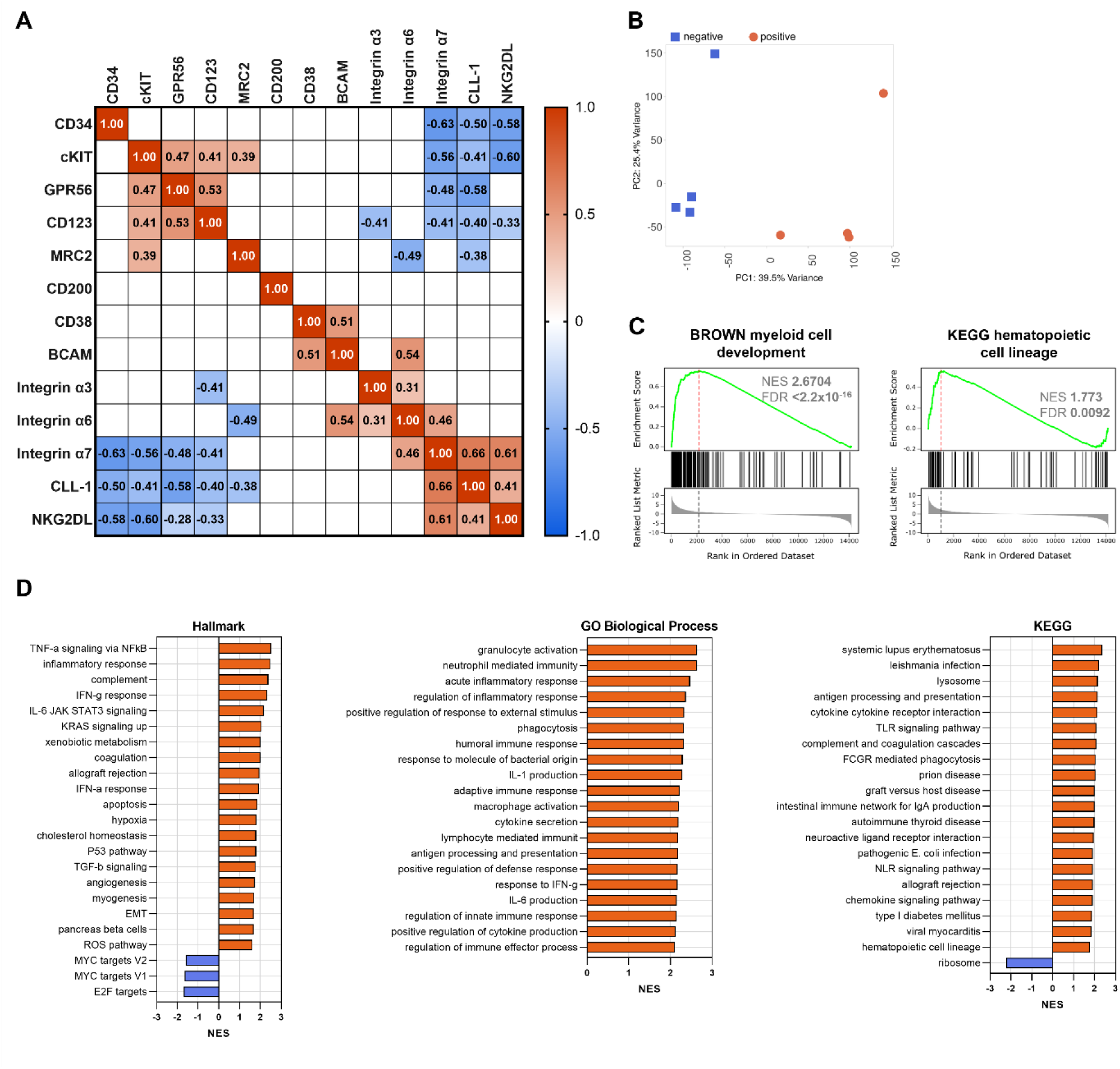
Integrin α7 expression is associated with differentiation. **(A)** Flow cytometric characterization of laminin receptors, LSC- and differentiation markers on primary AML cells. Integrin α7 expression negatively correlates with the expression of the LSC-markers CD34, cKIT, GPR56 and CD123. **(B)** Four primary AML samples were sorted for integrin α7 surface expression and RNA-sequencing was performed for both cell fractions. Principal Component Analysis (PCA) shows clustering of integrin α7+ and α7-samples despite a heterogeneous distribution within the groups. **(C)** GSEA shows enrichment of myeloid differentiation signatures in integrin α7+ samples. **(D)** GSEA using Hallmark, GOBP and KEGG from the Molecular Signature Database shows enrichment of molecular signatures associated with immune response and inflammation.

**Supplemental Figure S3.**
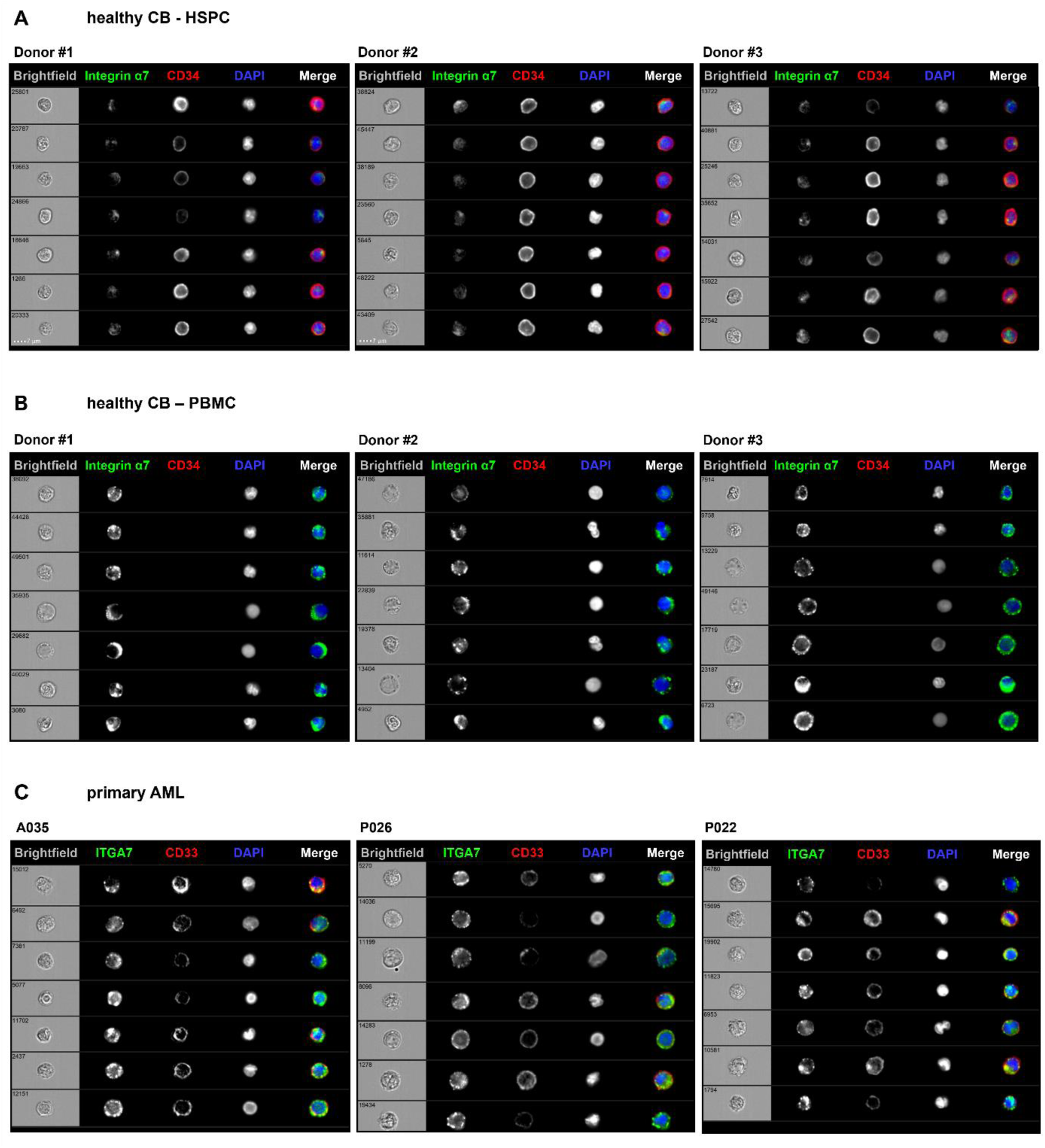
ImageStream analysis of subcellular integrin α7 localization in healthy HSPC, PBMC and primary AML cells. CB PBMC from healthy donors (n=3) and primary AML samples (n=3) were stained extracellularly and intracellularly for integrin α7. CD34 was stained on CB PBMC to distinguish HSPC. **(A, B)** In healthy CB PBMC, integrin α7 expression increases with the reduction in CD34 expression. While CD34+ HSPC predominantly show intracellular integrin α7 localization, PBMC show additional integrin α7 signals on the cell surface. **(C)** Primary AML cells express integrin α7 predominantly on the cell surface. Cells were acquired with Cytek Amnis ImageStream Mk II.

**Supplemental Figure S4.**
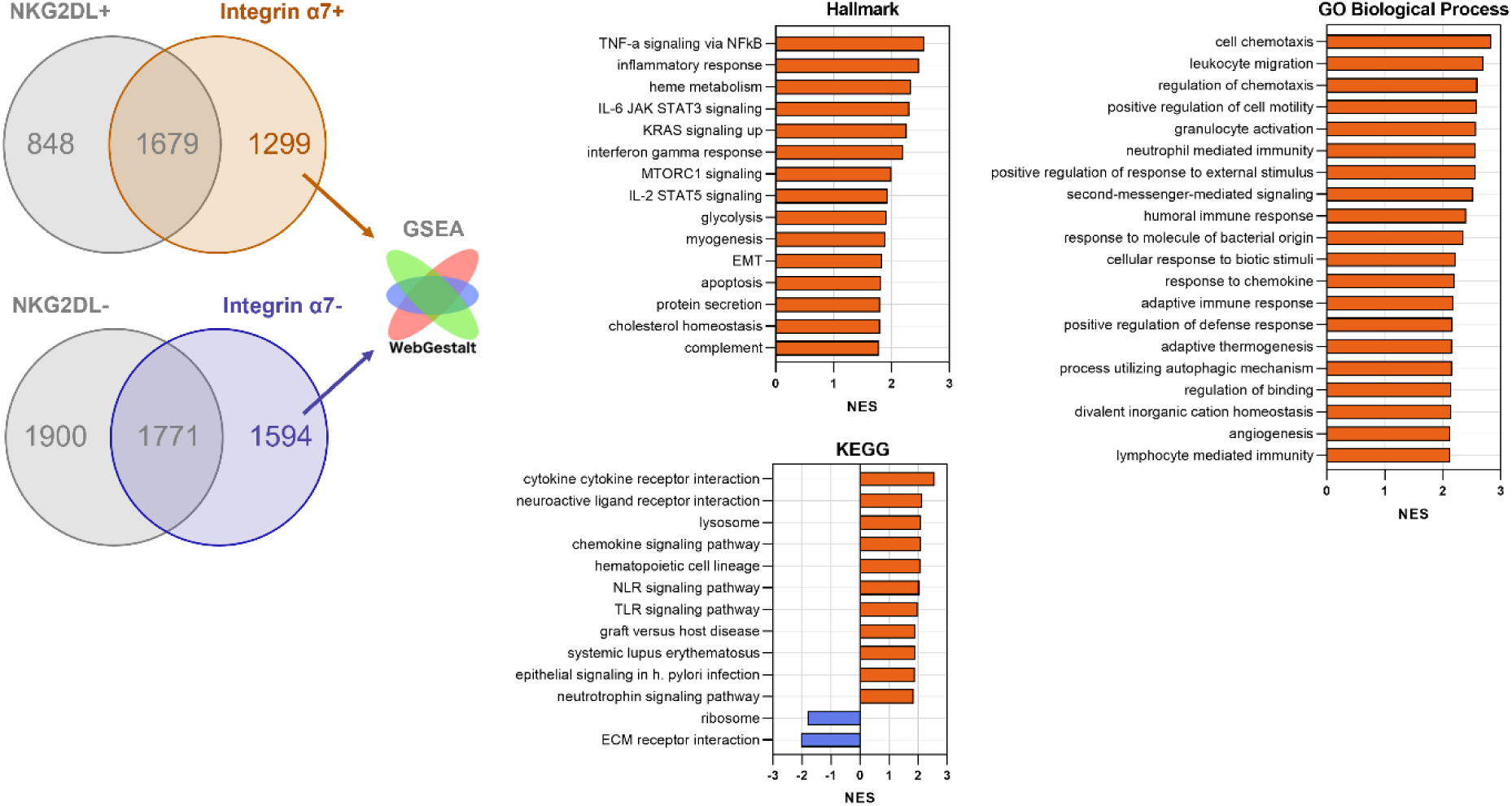
Comparison with the dataset GSE127959 reveals molecular signatures unique to integrin α7. Differentially expressed genes (logFC > 1) were compared with an LSC-dataset based on NKG2DL expression (logFC > 1). The genes unique to the integrin α7 dataset were used for GSEA. GSEA of genes unique to the integrin α7 dataset reveals enrichment of molecular signatures related to cell migration, angiogenesis and EMT.

**Supplemental Figure S5.**
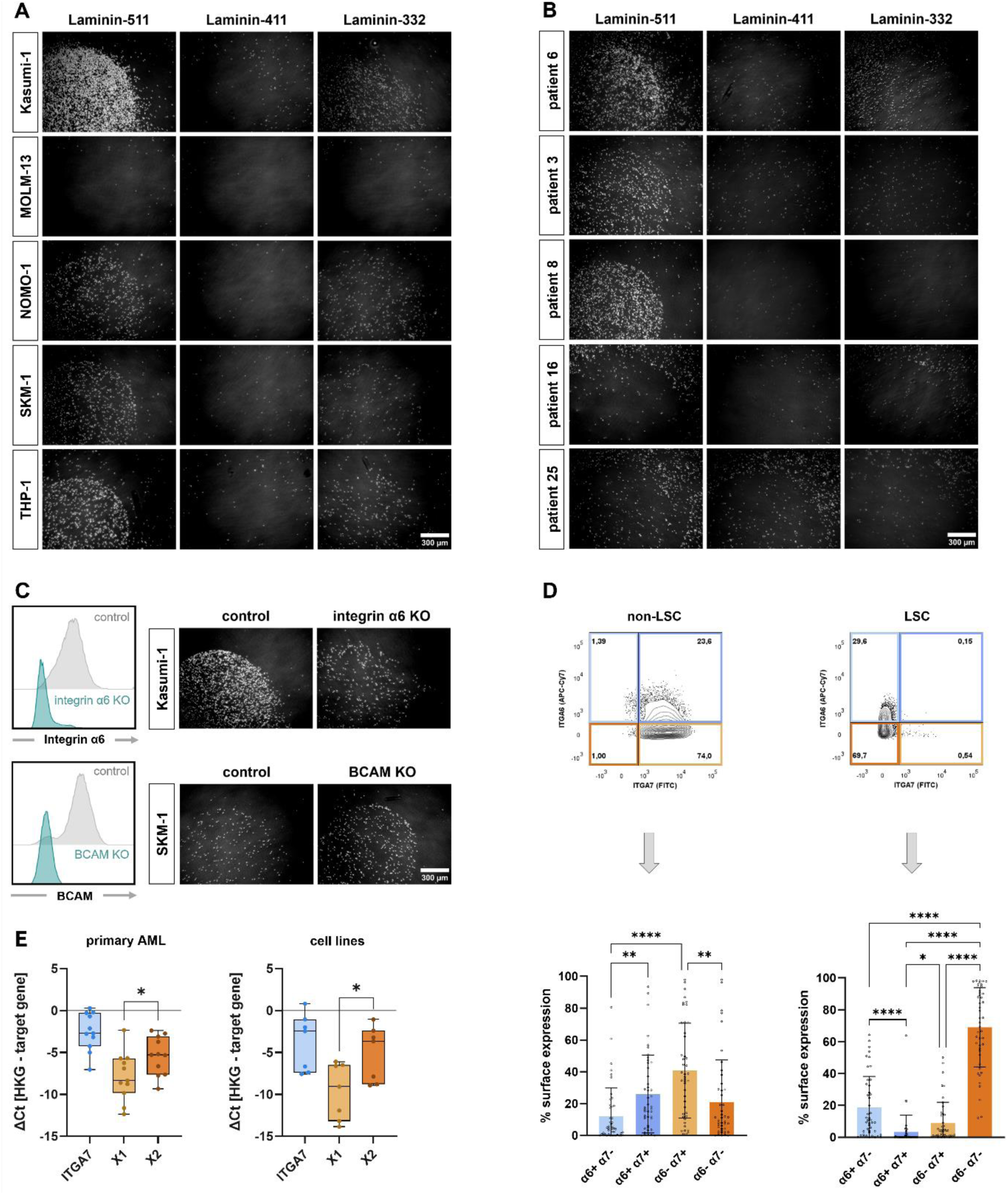
Laminin adhesion of AML cell lines and primary AML cells. Representative images of adhesion assays to different laminin isoform coatings. **(A)** Adhesion assays of AML cell lines show strong adhesion to laminin-511 and weak adhesion to laminin-332. All tested cell lines but MOLM-13 adhere to laminin-511. **(B)** Adhesion assays of primary AML cells show adhesion to laminin-511 in some samples, while other patient samples do not adhere. **(C)** Gene knockouts of integrin α6 and BCAM were generated in the AML cell lines Kasumi-1 and SKM-1 and the absence of surface expression of the respective marker was validated using flow cytometry. Deletion of integrin α6 reduces adhesion to laminin-511, but deletion of BCAM does not affect adhesion. **(D)** Flow cytometry data of primary AML samples analyzed for integrin α6 and α7 co-expression in non-LSC (NKG2DL+) and LSC populations (NKG2DL-). **(E)** QRT-PCR analysis of ITGA7 isoforms in primary AML cells (n=11) and AML cell lines (n=7). The isoform α7X2 is higher expressed than α7X1.

**Supplemental Figure S6.**
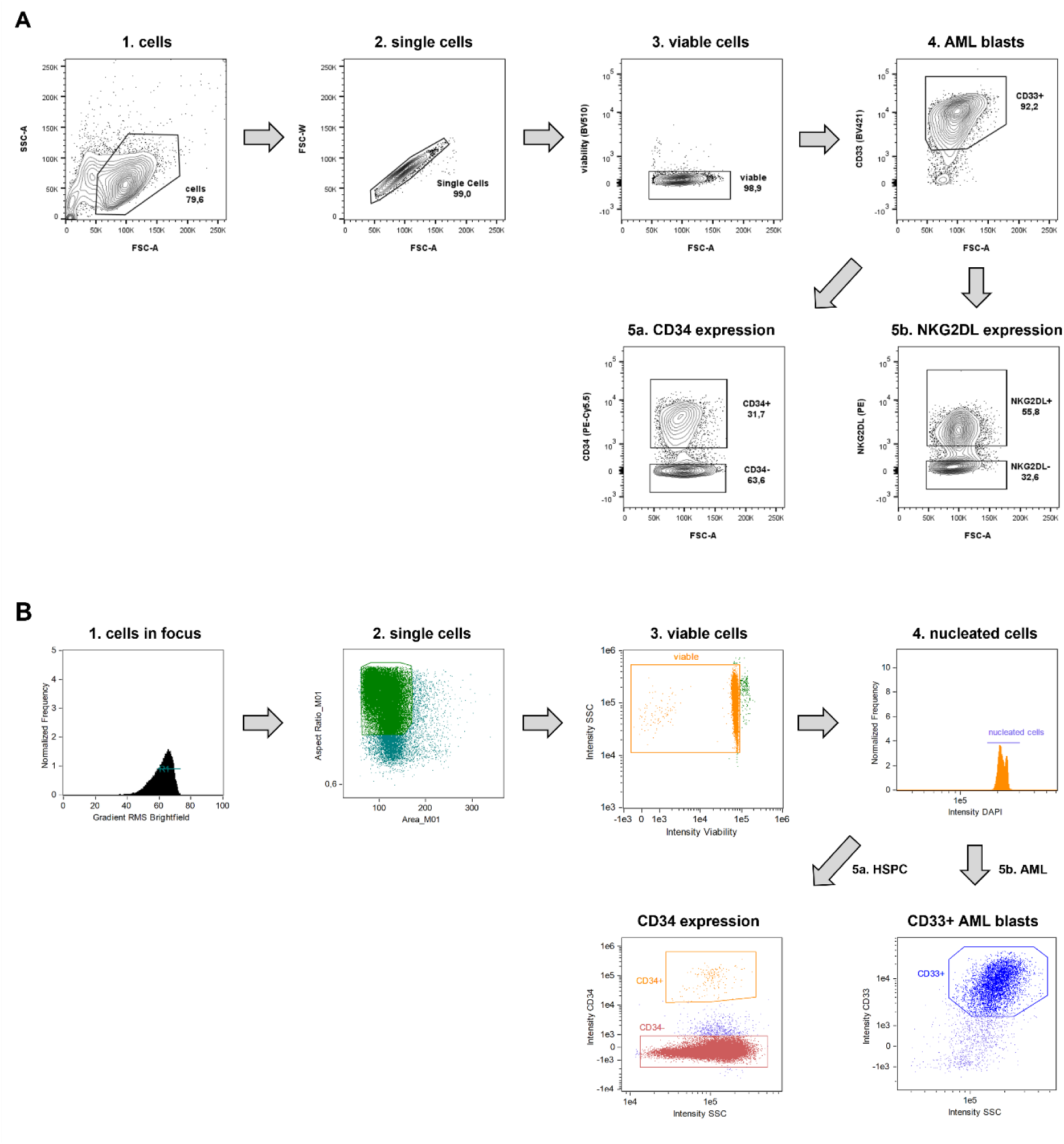
Gating strategy for flow cytometric and ImageStream analysis. **(A)** For flow cytometric analysis, single cells are defined via forward and sideward scatter. Dead cells are excluded and CD33 is used to define bulk AML blasts. CD34 and NKG2DL expression is used to distinguish LSC and non-LSC populations. **(B)** For ImageStream analysis, cells being in camera focus are selected. Single cells are defined by cell size and dead cells are excluded. DAPI intensity is used to define nucleated cells. In healthy samples, CD34 is used to distinguish HSPC from PBMC. In AML samples, CD33 is used to define bulk AML blasts.

